# Genomic and kinetic analysis of novel Nitrospinae enriched by cell sorting

**DOI:** 10.1101/2020.06.09.141952

**Authors:** Anna J. Mueller, Man-Young Jung, Cameron R. Strachan, Craig W. Herbold, Rasmus H. Kirkegaard, Michael Wagner, Holger Daims

## Abstract

Chemolithoautotrophic nitrite-oxidizing bacteria (NOB) are key players in global nitrogen and carbon cycling. Members of the phylum Nitrospinae are the most abundant, known NOB in the oceans. To date, only two closely affiliated Nitrospinae species have been isolated, which are only distantly related to the environmentally abundant uncultured Nitrospinae clades. Here, we applied live cell sorting, activity screening, and subcultivation on marine nitrite-oxidizing enrichments to obtain novel marine Nitrospinae. Two binary cultures were obtained, each containing one Nitrospinae strain and one alphaproteobacterial heterotroph. The Nitrospinae strains represent two new genera, and one strain is more closely related to environmentally abundant Nitrospinae than previously cultured NOB. With an apparent half-saturation constant of 8.7±2.5 µM, this strain has the highest affinity for nitrite among characterized marine NOB, while the other strain (16.2±1.6 µM) and *Nitrospina gracilis* (20.1±2.1 µM) displayed slightly lower nitrite affinities. The new strains and *N. gracilis* share core metabolic pathways for nitrite oxidation and CO_2_ fixation but differ remarkably in their genomic repertoires of terminal oxidases, use of organic N sources, alternative energy metabolisms, osmotic stress and phage defense. The new strains, tentatively named “*Candidatus* Nitrohelix vancouverensis” and “*Candidatus* Nitronauta litoralis”, shed light on the niche differentiation and potential ecological roles of Nitrospinae.

## Introduction

Bioavailable nitrogen is essential to all life on earth and the growth limiting factor in many ecosystems. With that said, in marine habitats, an excess in nitrogen availability due to anthropogenic pollution can cause dramatic effects such as coastal eutrophication and the formation of hypoxic dead zones [1]. The majority of fixed nitrogen in marine systems is composed of nitrate (NO_3_^-^) [2]. While microbially catalyzed, nitrite oxidation by chemolithoautotrophs is thought to be the most significant biological pathway in terms of nitrate production, surprisingly little is known about the underlying microbiology in the oceans. The known diversity of marine nitrite-oxidizing bacteria (NOB) consists of members of the phylum Nitrospinae and representatives of the genera *Nitrospira, Nitrococcus, Nitrobacter*, and *Nitrotoga* [3]. Among these, the Nitrospinae (recently proposed to be renamed to Nitrospinota [4]) are the most abundant NOB in the majority of the marine environments studied so far. They have been found to be present in a wide range of habitats including the free water column, oxygen minimum zones (OMZs), sediments, and deep-sea trenches [5-17]. Interestingly, nitrite oxidoreductase (NXR), the key enzyme for nitrite oxidation, was found to be highly abundant in metaproteomic studies of OMZs [11, 12] and high *in situ* nitrite oxidation rates were measured, although nitrite oxidation is considered to be an aerobic process and OMZs are strongly oxygen limited [5, 18, 19]. In addition to their importance in the nitrogen cycle, the Nitrospinae have been suggested to play a major role in dark ocean carbon fixation by contributing up to 15-45% of the fixed inorganic carbon in some environments [20]. The contribution of Nitrospinae to CO_2_ fixation is, however, still under debate [21, 22], partially due to a lack of cultured representative organisms.

To date, only two strains from the Nitrospinae have been isolated, both of which are members of the genus *Nitrospina* (*N. gracilis* and *N. watsonii*) and are closely related to each other (97.9% 16S rRNA identity) [23, 24]. Experimental and genomic evidence obtained thus far from these two cultured strains points to *Nitrospina* spp. being chemolithoautotrophic NOB and obligate aerobes [23–25]. However, single-cell genomic analyses of marine bacteria have revealed a much higher diversity of Nitrospinae. Accordingly, the phylum contains at least two additional major phylogenetic lineages, which are only distantly affiliated with the genus *Nitrospina* and are referred to as “Clade 1” (“*Candidatus* (*Ca*.) Nitromaritima”) and “Clade 2” [6, 20]. Intriguingly, these uncultured organisms are considerably more abundant in the environment than *Nitrospina* spp. [20]. They were predicted to be NOB based on the presence of *nxr* genes, but direct physiological evidence of this activity has been lacking. Various other NOB, such as *Nitrospira* members, have been shown to be able to utilize alternative energy metabolisms, including the oxidation of hydrogen or formate coupled to oxygen or nitrate reduction [26–28]. There is also some genomic and metagenomic evidence for putative alternative metabolisms in the Nitrospinae, such as hydrogen oxidation, sulfite oxidation, and nitrate reduction, which remain to be tested [6, 7, 20, 25]. Indeed, ecophysiological analyses of Nitrospinae outside the described genus *Nitrospina* are hampered by the lack of any cultivated strains that could systematically be characterized. The major cause of this deficiency is the recalcitrance of most NOB to cultivation and the time needed to isolate these organisms by traditional approaches [24, 29], which can take more than a decade.

We therefore employed an accelerated approach based on live cell sorting and obtained two binary co-cultures, each containing a novel Nitrospinae genus and a heterotrophic bacterium. Excitingly, one of the strains groups with the environmentally abundant but uncultured Nitrospinae clades 1 and 2 [20]. Here, we describe both strains, including genomic characterization and determination of their nitrite affinities, in order to expand our understanding of the diversity and ecophysiology of the Nitrospinae and provide new model organisms from this globally important phylum.

## Materials and Methods

### Sample collection, pre-enrichment and cultivation of NOB

Sandy coastal surface sediment (0-3cm) samples were taken at Burrard Inlet, Vancouver, Canada (49°16’22.1”N 123°11’32.5”W) in November 2016 and in Elba, Italy (42°43’48.1”N 10°09’23.2”E) in November 2016. Aliquots of the sediments were inoculated into 50 ml Schott glass bottles filled with 25 ml marine minimal medium at pH 7.4-7.6 containing 4.2 μl l^-1^ supplement solution (0.02g l^-1^ biotin, 0.02g l^-1^ folic acid, 0.10g l^-1^ pyridoxine HCL, 0.05g l^-1^ riboflavin, 0.05g l^-1^ nicotinic acid, 0.05g l^-1^ DL-pantothenic acid, 0.05g l^-1^ P-aminobenzoic acid, 2.00g l^-1^ choline chloride and 0.01g l^-1^ vitamin B_12_) and 0.5 mM NO_2_^-^, and incubated at 28°C in the dark without agitation [30]. The marine minimal medium was modified from the medium used for the cultivation of *N. gracilis* and *N. watsonii* by replacing natural seawater with 33.4 g l^- 1^ (Vancouver) and 39.9 g l^-1^ (Elba) red sea salts (Red Sea Aquaristic) dissolved in Milli-Q water [23–25]. A higher salinity was chosen for the Elba enrichment to reflect the higher salt concentrations in the Mediterranean Sea. The enrichments were regularly checked for nitrite consumption and nitrate formation by using nitrite/nitrate test stripes (Merckoquant, Merck). Upon consumption of nitrite, the cultures were replenished with 0.5 to 1 mM nitrite (Vancouver Nitrospinae) or up to 5 mM nitrite (Elba Nitrospinae and *N. gracilis*) (final concentrations). Once nitrite oxidation was consistently observed, aliquots of the cultures were transferred at a 1:10 ratio into fresh media. Unless stated otherwise, the Vancouver and Elba Nitrospinae were cultured as described above. *N. gracilis* was grown at the same red sea salt concentration as the Vancouver Nitrospinae. Growth of the Nitrospinae strains on solid media was tested on Marine Broth 2216 (Difco, BD) and the strain specific marine minimal media that were solidified with 15g l^-1^ agar.

### Cell sorting and activity screening

Cells were concentrated from 10 ml of the enrichment cultures by centrifugation (4500×g, 20 min, 28°C) using a swing-bucket rotor (Eppendorf) and resuspended in approximately 200 µl of the supernatant. Subsequently, the cells were sorted into sterile 96-well tissue culture plates (VWR, item no. 10062-900) by using the single cell sorting option at a pressure of 60 psi on a MoFlo Astrios Flow Cytometer (Beckman Coulter) equipped with a 70 µm jet in air nozzle. On the instrument, cells were visualized in the forward and side scatter channel and small, non-cell particles were excluded; otherwise no gating was applied. The wells of the microtiter plates contained 200 µl of sterile mineral salt medium amended with supplements (see cultivation details above) and 0.25 mM sodium nitrite. For the cultivation of the Vancouver sourced Nitrospinae strain, the sorting medium additionally contained 0.1 mM sodium pyruvate to alleviate oxidative stress [31]. The 96-well microtiter plates were placed in closed plastic bags to prevent evaporation, incubated at 28°C in the dark without agitation and 10 µl aliquots of each well were regularly checked for nitrite consumption with the Griess assay [32]. Selected wells, which showed nitrite consumption, were gradually scaled up by transferring the cultures into larger volume microtiter plates and doubling the culture volume after each round of nitrite consumption up to 3.2 ml and then to a final volume of 25 or 50 ml in Schott bottles. The cultures were further cultivated as described for the enrichments (see above). Once sufficient biomass was available, the enriched nitrite-oxidizing organisms were provisionally identified by full-length 16S rRNA gene amplification and Sanger sequencing.

The morphology of the cells was visualized via scanning electron microscopy and catalyzed reporter deposition fluorescence *in situ* hybridization (CARD-FISH) with the Nitrospinae specific 16S rRNA-targeted probe Ntspn759 [21] (Supplemental Materials and Methods).

### Genomic and phylogenetic analyses

DNA was extracted from the cultures and complete genomes were obtained through Illumina and Nanopore sequence co-assemblies. A 16S rRNA gene phylogenetic tree was calculated and the genomes were annotated on the MicroScope platform (MAGE Workflow version: 1.8 [33] (Supplemental Materials and Methods).

Publicly available Nitrospinae genomes, including metagenome assembled genomes (MAGs) and single-cell amplified genomes (SAGs), were retrieved from NCBI and from the JGI genome portal (see Table S1 for details). Completeness and contamination of the genomes was assessed by CheckM (v. 1.0.18) (Table S1), and phylogenetic analyses were conducted on genomes that were more than 80% complete and less than 10% contaminated [34]. A phylogenetic tree was calculated with an alignment of concatenated conserved bacterial marker proteins, made with the GTDB (Genome Taxonomy Database) toolkit (v. 0.3.2), using IQ-TREE (v. 1.6.11, model: LG+F+R5 as chosen by automatic model selection and 1000 ultrafast bootstrap runs) [4, 35–37]. The genomes of *Nitrospira moscoviensis, G. metallireducens*, and *D. multivorans* served as outgroup. Clades within the Nitrospinae were depicted as they have been previously described in the literature [6, 20]. Average amino acid identity (AAI) and whole-genome nucleotide identity (gANI) were calculated as described elsewhere [38–40]and visualized using R (v. 3.6.1) with the R package tidyverse (v. 1.3.0) [41]. The GTDB-TK output (Table S1) was used to delineate phylogenetic affiliations beyond the genus level [4].

The raw reads and genomes of the novel Nitrospinae strains were submitted to NCBI under the Bioproject accession PRJNA602816 and the Biosamples SAMN13976148 (“*Ca*. Nitrohelix vancouverensis” VA) and SAMN13976151 (“*Ca*. Nitronauta litoralis” EB). The genomes are further available on the MicroScope platform under the accession NTSPN23.2 (“*Ca*. Nitrohelix vancouverensis” VA) and NTSPN 3.2 (“*Ca*. Nitronauta litoralis” EB).

### Nitrite oxidation kinetics

Nitrite oxidation kinetics of the Nitrospinae co-cultures and *N. gracilis* were quantified by measuring nitrite-dependent oxygen consumption in a microrespirometry (MR) system (Unisense) as described in detail by Kits *et al*. [42]. To concentrate the biomass for MR, cells were collected from 100 to 300 ml of culture with Amicon Ultra-15 tubes (Millipore) by centrifugation (4500×g, 10 min, 28°C) using a swing-bucket rotor (Eppendorf). Concentrated biomass was washed and resuspended in nitrite free growth medium for the MR experiments. The media did not contain supplements to prevent oxygen respiration due to the degradation of organic compounds by the co-enriched, heterotrophic bacteria during the MR experiment. Culture biomass was incubated in a water bath set to the experimental temperature prior to being transferred to a 2 ml glass MR chamber with a stir bar. All MR experiments were performed with 300 rpm stirring at 28°C. Small culture volumes (∼10 µl) were taken before and immediately after nitrite injection, and ∼5 min after nitrite depletion for nitrite/nitrate measurements to confirm stoichiometric conversion of oxygen, nitrite and nitrate. Protein concentrations were determined with the Pierce BCA protein assay (ThermoScientific) and cell abundances by qPCR (Supplementary Methods). All experiments were replicated three times or more, using at least two different cultures on different days (see Results and Discussion for total biological and technical replicates per strain). For one replicate of each strain, images of the cells stained with DAPI were taken before and after the MR measurements by using an epifluorescence microscope (Zeiss). These images were used to check whether the cells in the MR chamber had formed aggregates, which could cause oxygen diffusion limitations and thus affect the measured respiration rates and the inferred nitrite oxidation kinetics. No aggregates were observed.

## Results and Discussion

### Cultivation of novel Nitrospinae representatives

For the initial enrichment of marine NOB, nitrite-containing mineral media were inoculated with coastal surface sediment samples taken in Vancouver, Canada, and on Elba, Italy. Within 4 weeks of incubation, nitrite oxidation to nitrate was detected in the cultures and this activity continued to be observed after subsequent replenishment of nitrite and transfers of culture aliquots into fresh medium. Usually, the further purification of NOB from accompanying organisms in the enrichment cultures is hindered by the very slow growth and inability of most NOB (including all cultured Nitrospinae) to grow on solid media. To expedite the purification of Nitrospinae strains from our initial enrichments, which was previously a laborious and lengthy process [24], we developed a method for the physical separation, activity-based identification of NOB, and subcultivation in 96-well microtiter plates. This method uses random, non-fluorescent, single-cell sorting using a fluorescence activated single cell sorting (FACS) instrument paired with a nitrite consumption activity screen (Fig. S1a). Thus it differs from a previously reported FACS isolation approach for *Nitrospira* NOB from activated sludge, where the NOB were targeted based on their known cell cluster size and shape, which had been determined by *Nitrospira*-specific rRNA-targeted FISH analysis before FACS was performed [43]. Our method does not rely on prior knowledge of the identity and morphology of the NOB [43], but is solely based on the detection of nitrite oxidation after sorting. Still, it might allow for a flexible selection of the sorted cell morphologies by adjusting the gating parameters. While the previous method facilitated the isolation of already known NOB, the approach used in our study was designed for the discovery of novel nitrite oxidizers that grow under the given conditions. It may also be suitable for the isolation of other microorganisms that can be efficiently sorted (i.e., grow in suspension or that can be suspended by sonication or other methods) and that perform a specific metabolism of interest, which is detectable by a colorimetric, fluorimetric, or other high- throughput assay. Examples include previously performed, high-throughput enzyme discoveries and the isolation of microalgae [44–46]. After sorting cells from the initial Vancouver and Elba enrichment cultures into one 96-well plate each, several wells showed nitrite-oxidizing activity within 3-4 weeks. Cells from three active wells from the Elba enrichment and 4 for the Vancouver enrichment (>4 were active) were progressively transferred into larger culture volumes. This procedure led to the separate enrichment of two different Nitrospinae, one from each of the initial enrichments, that were identified by 16S rRNA gene sequencing (Fig. S1b): strain VA (Vancouver) and strain EB (Elba). Interestingly, despite attempted single-cell sorting and various dilution to extinction attempts, an axenic culture could not be established for either of the two obtained Nitrospinae strains. Rather, these cultures represent binary co-cultures each containing one Nitrospinae strain, one alphaproteobacterial strain, and no other detectable microorganisms. Overall, using this single-cell sorting and screening method, we were able to obtain the two binary co-cultures from the environmental samples in approximately 10 months.

### Co-enrichment with Alphaproteobacteria

Pure cultures of the two co-enriched heterotrophic strains were obtained on Marine Broth Agar. Subsequent 16S rRNA gene analysis of these isolates showed that the two Nitrospinae strains had been co-cultured with members of two distinct genera within the Alphaproteobacteria. Strain VA was co-cultured with a bacterium most closely related to *Stappia stellulata* strain (NR_113809.1, 99.58% 16S rRNA gene sequence identity), whereas the EB strain culture contained a *Maritimibacter alkaliphilus* strain (NR_044015.1, 100% 16S rRNA gene sequence identity). Both these species have previously been isolated from marine environments and have been described as alkaliphilic chemoorganoheterotrophs that can use a wide variety of simple and complex organic substrates [47, 48]. In our cultures, which were only provided with nitrite for growth, the alphaproteobacterial strains may have lived off simple organic compounds that were excreted by the autotrophic Nitrospinae. Since the NOB could not be grown separately from the heterotrophs, it is tempting to speculate that the Nitrospinae strains also benefitted, for example, from reactive oxygen species (ROS) protection by the heterotrophs. Superoxide dismutase and catalase genes are present in the genomes of both co-cultured alphaproteobacteria, and other isolates of both species are catalase positive [47, 48]. Similar interactions have already been observed in marine autotroph-heterotroph co-cultures, including other nitrifiers [49–51].

### Phylogeny of the novel Nitrospinae

Closed genomes were reconstructed for both Nitrospinae strains by co-assembling Illumina and Nanopore sequencing data. Since the genome of *N. gracilis* was nearly completely sequenced but not closed [25] (Table S2), the obtained genomes represent the first closed genomes from Nitrospinae. With a length of >3.9 Mbp, Nitrospinae strain EB has a larger genome than the other cultured Nitrospinae strains sequenced to date (Table S2).

Phylogenetic analysis of the 16S rRNA genes (Fig. S1b) showed that the Nitrospinae strains VA and EB are only distantly related to *N. gracilis* (93% sequence identity with EB and 91% with VA), *N. watsonii* (92% sequence identity with EB and 90% with VA), and to each other (90% sequence identity). Since public databases contained only few 16S rRNA gene sequences closely related to strains VA and EB (Fig. S1b), these strains seem to represent a yet underexplored diversity of Nitrospinae. A phylogenetic tree using conserved concatenated marker proteins from all available Nitrospinae genomes that are >80% complete and <10% contaminated, including MAGs and SAGs, further confirmed that the strains obtained here represent novel lineages (Fig. 1). In particular, strain VA was more closely related with the Clade 1 and 2 Nitrospinae than *N. gracilis*, although it was clearly not a member of either group. In the concatenated marker tree, it belonged to a third well-resolved clade with additional MAGs. The exact branching order between Clade 1, 2, and this third clade remained unresolved (Fig. 1, Fig. S1b). The novelty of the two cultured Nitrospinae strains was corroborated by genome average nucleotide identity (gANI) (Fig. S2) and average amino acid identity (AAI) (Fig. 2) analyses. Since strain VA, strain EB, and most of the Nitrospinae members in the dataset are quite distantly related (below the 96.5% species level cut-off [39]), gANI (Fig. S2) was not suitable to further resolve their taxonomic grouping. However, AAI analysis revealed a high genus-level diversity within the Nitrospinae, comprising 15 distinct genera according to a proposed 60% genus level cut-off [52]. Both strains from this study, VA and EB, represent a new genus with 55% and 58% AAI to *N. gracilis*, respectively, and 53% AAI to each other (Fig. 2). The AAI analysis also suggested that each of the environmental clades 1 (“*Ca*. Nitromaritima”) and 2 represent separate genera, with the lowest AAI values within each clade being >70%, and that these genera are distinct from the genus containing strain VA (Fig. 2). In order to make taxonomic inferences beyond the genus level, the GTDB-TK tool [38] was applied to the Nitrospinae genome dataset. GTDB-TK confirmed that strains VA and EB belonged to the Nitrospinaceae family but, in agreement with the AAI analysis, did not assign them to a genus (Tab. S2). While we showed that nitrite oxidation is spread among phylogenetically distant members within the Nitrospinaceae, no nitrite oxidation phenotype has been observed yet for the Nitrospinae members that do not belong to the Nitrospinaceae family.

**Figure 1.**
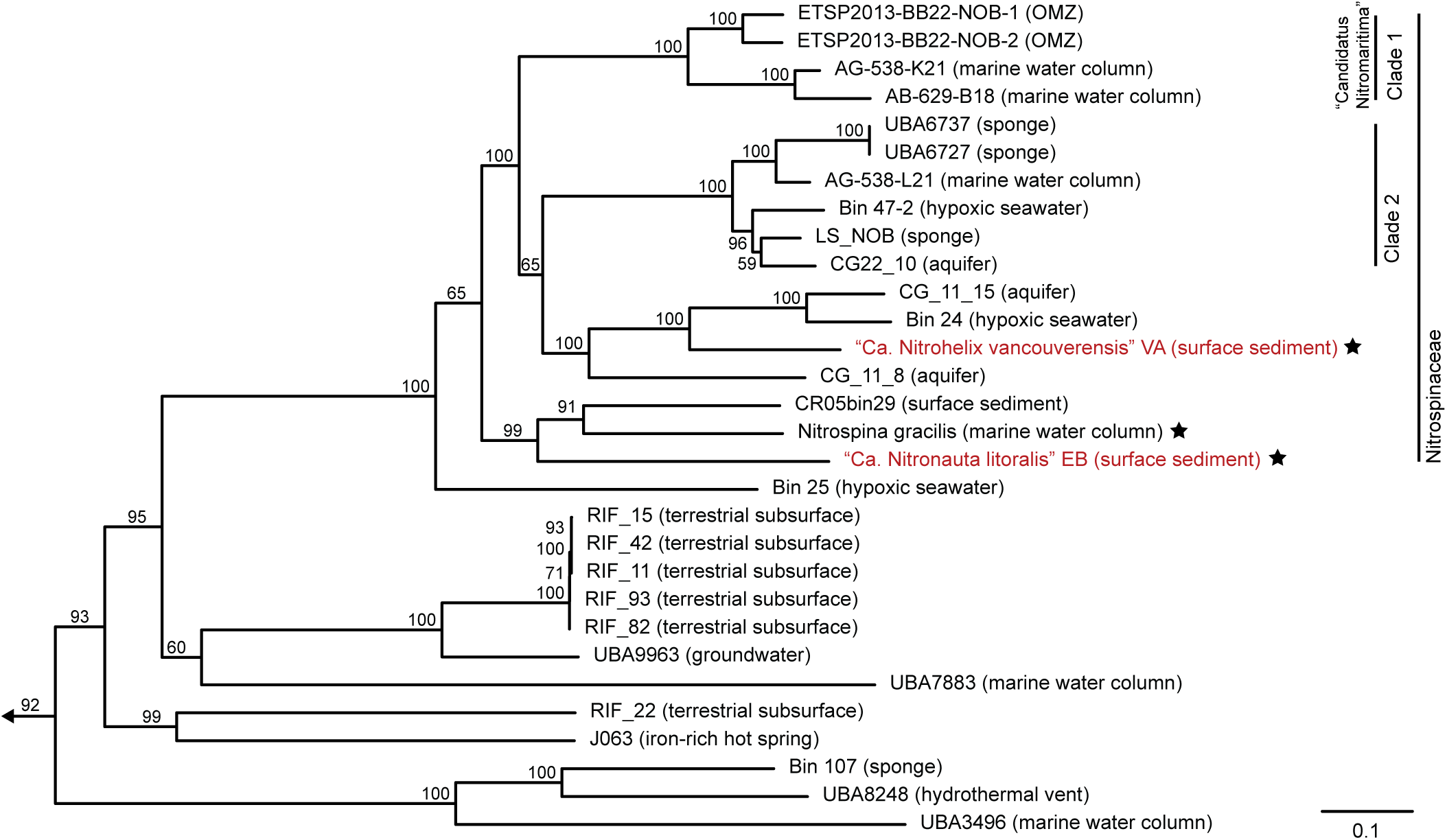
Concatenated marker gene tree of the Nitrospinae. The maximum likelihood tree shows the phylogenetic positions of the two newly cultured strains (highlighted in red) within the phylum Nitrospinae. The tree was calculated from a concatenated alignment of 120 conserved bacterial marker proteins. Details of the Nitrospinae genomes, which were used to calculate this tree, are listed in Table S1. Numbers at branches indicate ultrafast bootstrap (n=1,000) support. *Nitrospira moscoviensis, Desulfococcus multivorans*, and *Geobacter metallireducens* were used as an outgroup. Cultured organisms are marked with an asterisk. Clades of the Nitrospinae are indicated as proposed elsewhere [6] and the family Nitrospinaceae is indicated as determined by GTDB-TK. The scale bar indicates 0.1 estimated substitutions per residue. The sample source is indicated in parentheses.

**Figure 2.**
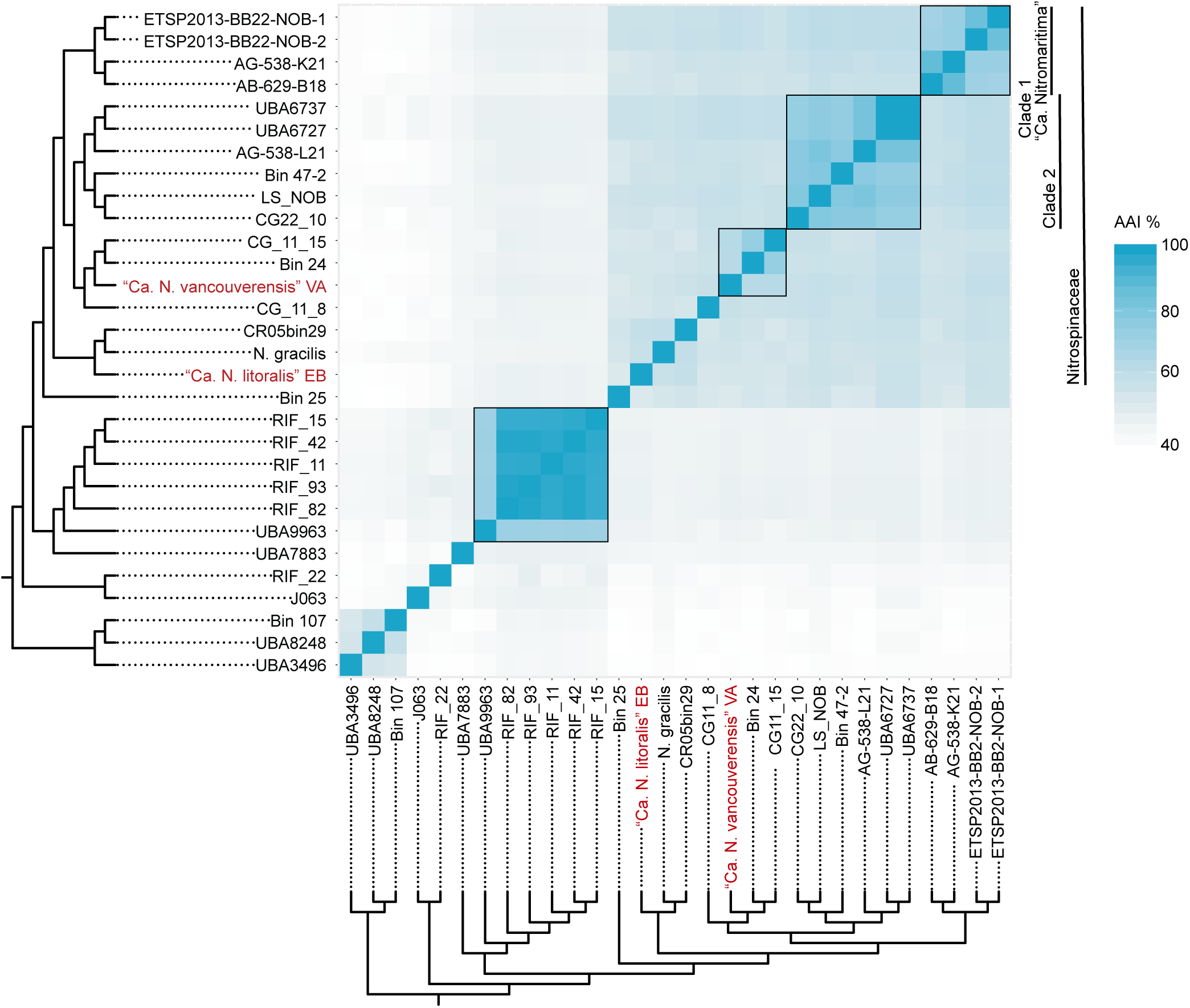
Average amino acid identity (AAI) analysis of the Nitrospinae. Pairwise AAI values were calculated for the same set of Nitrospinae genomes that was used to reconstruct the phylogenetic tree in Fig. 1. The same tree is used here to annotate the heatmap. The two newly cultured strains are highlighted in red. Clades of the Nitrospinae are indicated as proposed elsewhere [6]. The boxes indicate the genus boundary at 60% identity.

Taken together, the phylogenetic, ANI, AAI, and GTDB-TK analyses revealed a high genus-level diversity within the Nitrospinae, the vast majority remaining uncultured and poorly characterized. Among all cultured Nitrospinae members, strain VA is most closely related with the uncultured but environmentally abundant clades 1 and 2 [6, 7, 20, 21].

### Cell morphology

The morphologies of the new Nitrospinae strains were visualized using scanning electron microscopy (SEM), and by 16S rRNA-targeted CARD-FISH using a Nitrospinae-specific oligonucleotide probe (Ntspn759) [21]. Strain VA cells were helically shaped rods (Fig. 3a, d, g), whereas strain EB appeared to be short, slightly curved rods (Fig. 3b, e, h). Interestingly, neither of the two strains displayed the long, slender, rod-shaped morphology that was mostly observed for the previously isolated *Nitrospina* species, *N. gracilis* (Fig. 3c, f, i) [23]. The shape of strain VA (Fig. 3a, d, g) rather resembled *Nitrospira* species [53]. The short, slightly curved rod morphology of strain EB (Fig. 3b, e, h) resembled the compact cell shape reported for aging cultures of *N. watsonii* [24] and environmental Nitrospinae [5]. While the helical shape of strain VA could be clearly distinguished from the co-cultured *Stappia* sp. using Nitrospinae-specific FISH (Fig. 3g) and SEM of the *Stappia*-like isolate (Fig. S3), assigning a morphology to strain EB was slightly more difficult due a more similar morphotype of the co-cultured *Maritimibacter*- like bacterium (Fig. 3e and Fig. S3b). According to SEM, the isolated *Maritimibacter* had a coccoid morphology (Fig. S3) similar to the slightly smaller coccoid cells that were observed in the active co-culture with strain EB (Fig. 3e). Therefore, we assume that the slightly larger, curved rods in the SEM pictures (Fig. 3b, e) were Nitrospinae strain EB cells. However, the previously described morphological variability suggests that the cell shape of Nitrospinae is influenced by the growth stage and environment [23, 24]. Thus, morphology would be of limited use as the sole criterion to differentiate Nitrospinae strains from each other and from other organisms. We propose the name “*Candidatus* Nitrohelix vancouverensis” VA for strain VA based on its observed morphology and isolation source, and “*Candidatus* Nitronauta litoralis” EB for strain EB based on its isolation source.

**Figure 3.**
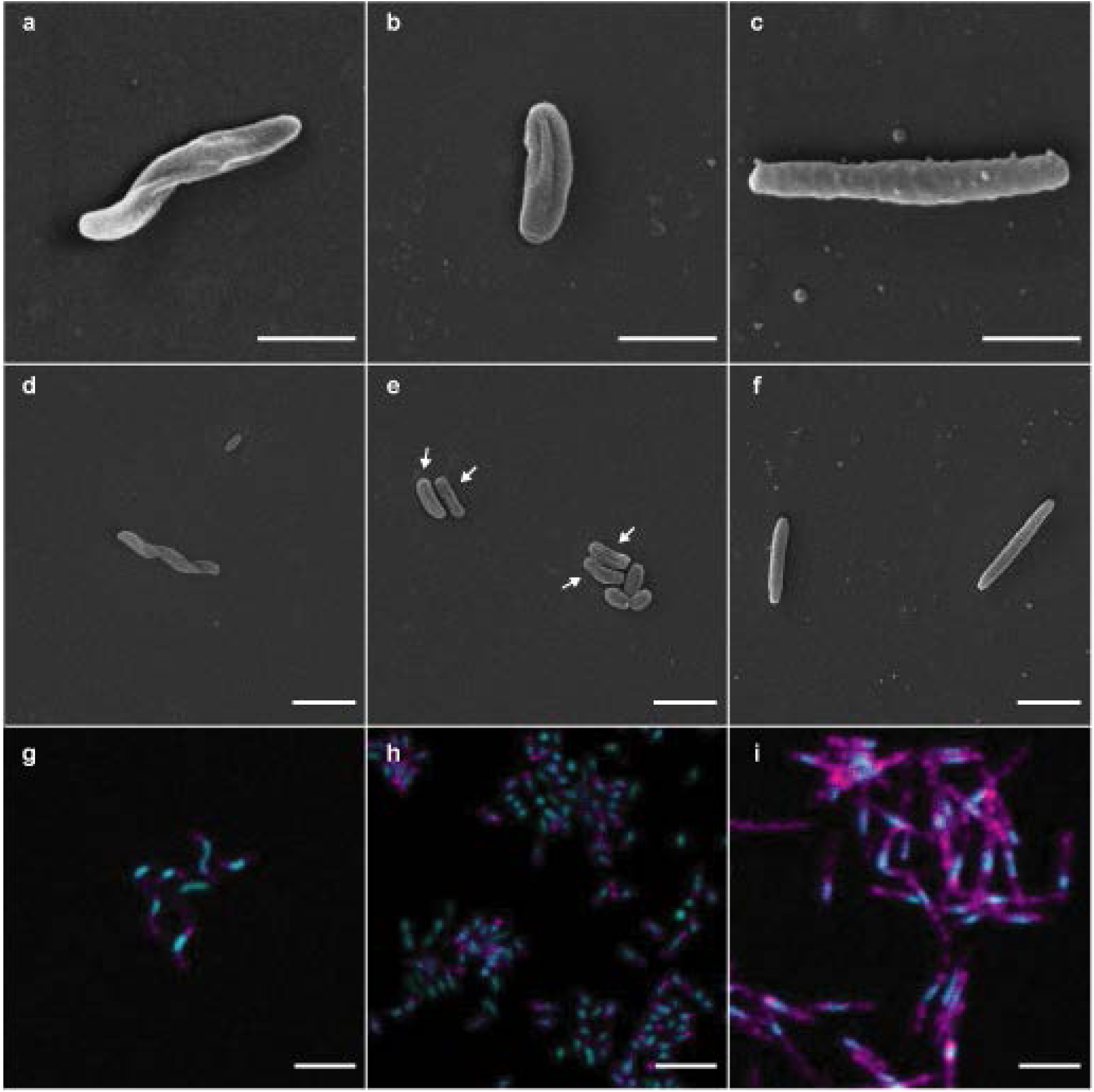
Scanning electron microscopy (SEM) and CARD-FISH images of the two newly cultured Nitrospinae strains and *N. gracilis*. **a, d, g** “*Ca*. Nitrohelix vancouverensis” strain VA. **b, e, h** “*Ca*. Nitronauta litoralis” strain EB. **c, f, i** *N. gracilis*. The short rod in **d** is a cell of the accompanying heterotroph *Stappia* sp. in the enrichment. **e** Shows slightly longer and slightly curved rods (strain EB, arrows) and shorter rods, which are cells of the accompanying heterotroph *M. alkaliphilus* in the enrichment. The scale bars in **a-c** depict 1 µm; all other scale bars depict 2 µm. **g-i** The 16S rRNA-targeted probe Ntspn759 was used to detect Nitrospinae cells (magenta), and total nucleic acids were stained with SYBR Green (cyan).

### Nitrite oxidation: activity and kinetics

Both Nitrospinae strains oxidized nitrite stoichiometrically to nitrate (Fig. S4). Exponential growth of “*Ca*. N. vancouverensis” correlated with the consumption of nitrite (Fig. S4a), whereas “*Ca*. N. litoralis” did not enter the exponential growth phase during the incubation period (Fig. S4b). During the experiment, the relative abundance of “*Ca*. N. vancouverensis” compared to the co-cultured *Stappia* sp. increased pronouncedly from 6 to 75% (Fig. S4a). The relative abundance of “*Ca*. N. litoralis” compared to the *Maritimibacter sp*. could not be reliably determined during the incubation experiment (Fig. S4b), but measurements taken after the MR experiments (see below) showed that the relative abundance of “*Ca*. N. litoralis” ranged from approximately 56 to 99%. Thus, the quantitative composition of each co-culture appeared to fluctuate and likely depended strongly on the availability of nitrite as the substrate for the NOB strains.

The kinetics of nitrite oxidation of *N. gracilis* and the two novel Nitrospinae strains were assessed by measuring the nitrite-dependent oxygen consumption in MR experiments. For all three NOB, nitrite oxidation followed Michaelis-Menten kinetics (Fig. S5). The stoichiometry of NO_2_^-^ and O_2_ consumption was always close to 1:0.5 (*N. gracilis*: mean=1:0.49; s.d.=0.02; *n*=8; “*Ca*. N. vancouverensis”: mean=1:0.51; s.d.=0.01; *n*=4; “*Ca*. N. litoralis”: mean=1:0.51; s.d.=0.02; *n*=5). This ratio was expected for NOB [54] and indicates that O_2_ consumption by the co-enriched heterotrophs was only minor during the relatively short MR experiments (maximum 1 h) and did not affect the kinetic analysis of “*Ca*. N. vancouverensis” and “*Ca*. N. litoralis”. The apparent half-saturation constant, *K*_m(app)_, of *N. gracilis* was determined to be 20.1 µM NO_2_^-^ (s.d.=2.1, *n*=8) and the maximum reaction rate, V_*max*_, to be 41.4 µmol NO_2_^-^ mg protein^-1^ hour^-1^ (s.d.=9.4, *n*=6), which is highly similar to the previously reported *K*_m(app)_ and V_*max*_ of the closely related *N. watsonii* (18.7±2.1 µM NO_2_^-^ and 36.8 µmol NO_2_^-^ mg protein^-1^ hour^-1^) [55]. The *K*_m(app)_ measured for “*Ca*. N. litoralis” was 16.2 µM NO_2_^-^ (s.d.=1.6, *n*=7) and thus resembled the values of *N. gracilis* and *N. watsonii*. In contrast, with a *K*_m(app)_ of 8.7 µM NO_2_^-^ (s.d.=2.5, *n*=3), “*Ca*. N. vancouverensis” showed a higher affinity for nitrite that was comparable with non-marine *Nitrospira* members (*K*_m(app)_=6 to 9 µM NO_2_^-^), which have been the cultured NOB with the highest nitrite affinity known so far [54, 56]. Indeed, strain VA turned out to have the lowest *K*_m(app)_ of all hitherto analyzed marine NOB in culture (Fig. 4). Interestingly, among all cultured marine NOB, “*Ca*. N. vancouverensis” is also most closely related to the Nitrospinae clades 1 and 2 that are abundant in oligotrophic waters [20] (see above). However, its *K*_m(app)_ is still 1-2 orders of magnitude higher than the *K*_*m(*app)_ values of nitrite oxidation reported for environmental samples from an OMZ (0.254±0.161 μM NO_2_^-^) and South China Sea waters (0.03 to 0.5 µM) [22, 57]. The very high nitrite affinity observed with these samples might be explained by the presence of uncharacterized nitrite oxidizers, whose nitrite affinity exceeds that of all cultured NOB. However, it remains to be tested whether known NOB can persist under extremely low *in situ* nitrite concentrations. For example, the half-saturation constant of growth for ammonia oxidizing bacteria spans several orders of magnitude under different temperatures [58]. Strongly different substrate affinities have also been observed for *Nitrobacter winogradskyi* and *Escherichia coli* under oligotrophic versus copiotrophic growth conditions [59, 60]. Systematic assessments of the kinetic plasticity of NOB under different conditions are still pending, mainly because the production of sufficient biomass of NOB isolates has been a major obstacle for such studies.

**Figure 4.**
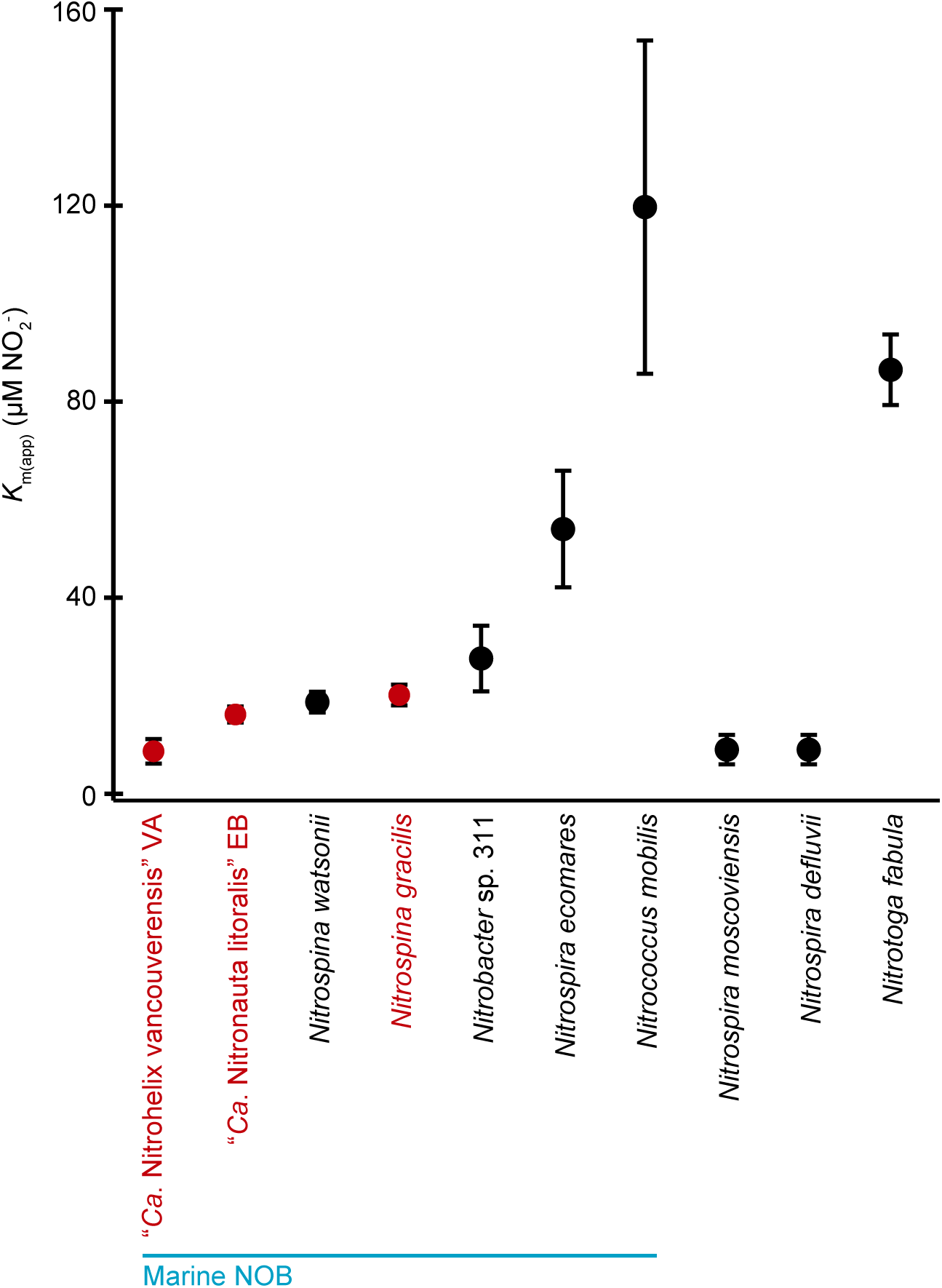
Comparison of the whole-cell apparent half-saturation constants (*K*_m(app)_) for nitrite between marine and non-marine NOB. The *K*_m(app)_ values measured in this study (highlighted in red) are the mean from all biological replicates (n=3 for “*Ca*. Nitrohelix vancouverensis” VA; n=5 for “*Ca*. Nitronauta litoralis” EB; n=8 for *N. gracilis*). The other *K*_m(app)_ values were retrieved from previous studies [54, 55, 81].

### General genomic features of cultured Nitrospinae

A pan-genomic analysis of the two novel cultured Nitrospinae strains and *N. gracilis* showed that these three organisms share a core genome of 1347 proteins, which have at least 50% amino acid sequence identity over 80% of the alignment (Fig. S6). The core genome included universally highly conserved bacterial genes, such as those coding for ribosomal proteins, translational elongation factors and the DNA replication machinery, as well as the genes for the core metabolism of chemolithoautotrophic NOB. Interestingly, among the shared conserved genes we also found highly conserved glutaredoxins, thioredoxin reductases, and peroxidases (>80% amino acid identity among the respective homologs). Like *N. gracilis* [25], “*Ca*. N. vancouverensis” and “*Ca*. N. litoralis” lack the canonical defense systems of aerobic organisms against oxidative stress, catalase and superoxide dismutase. While the aforementioned core genes could thus be essential for the detoxification of peroxides in all three organisms [61, 62], it remains a mystery how Nitrospinae deal with superoxide radicals [25]. Each of the strains encode a number of unique proteins (Fig. S6), many of which are phage related, corroborating a recently proposed hypothesis predicting extensive phage predation on Nitrospinae [21] (Supplemental Results and Discussion, Fig. S8). Yet, the majority of the variable genome content is still functionally uncharacterized. However, a few genes of the variable genome have known functions and might be important for niche adaptations. In the following sections, we address these genes as well as the shared core metabolism of the three analyzed Nitrospinae.

### Nitrite oxidation and respiration

Among the highly conserved proteins are the three known subunits of a periplasmic nitrite oxidoreductase, NxrABC. Details of the predicted subunit composition and cofactors of the NXR of Nitrospinae, which is closely related to the NXR of *Nitrospira*, have been described elsewhere [3, 25, 63]. Briefly, all three Nitrospinae strains possess two genomic copies of the substrate-binding subunit NxrA, two or three (only “*Ca*. N. vancouverensis”) copies of the electron- channeling subunit NxrB (Fig. S7), and several copies of putative NxrC subunits, which may transfer electrons from NXR to a downstream acceptor in the electron transport chain (Table S4). Homologs to all of the different putative NxrC subunits of *N. gracili*s [25] were also found in “*Ca*. N. vancouverensis” and “*Ca*. N. litoralis” (Table S4), but it remains to be determined whether all of these proteins are functionally involved in nitrite oxidation.

The respiratory electron transport chain of NOB is short, as electrons derived from nitrite are directly transferred, via *a*- or *c*-type cytochromes, to the terminal oxidase (complex IV) [25, 63, 64]. *N. gracilis* carries a *cbb*_*3*_-type high affinity heme-copper cyt. *c* oxidase (HCO) [25], whereas both “*Ca*. N. vancouverensis” and “*Ca*. N. litoralis” lack any canonical HCO. However, all three organisms encode highly conserved, putative “*bd*-like oxidases” [25]. These proteins, which also occur in all *Nitrospira* genomes, are phylogenetically related with but clearly distinct from the canonical cyt. *bd*-type quinol oxidases [63]. Interestingly, one variant of the *bd*-like oxidases from *Nitrospira* contains all conserved amino acid residues for heme and Cu binding in HCOs [65], indicating that this enzyme could be a novel cyt. *c*-oxidizing HCO [63]. The *bd*-like oxidases of *N. gracilis*, “*Ca*. N. vancouverensis”, and “*Ca*. N. litoralis” have most of these conserved residues; however, one of the three histidine ligands of the Cu_B_ is replaced with a glutamine, and the histidine ligand of the high-spin heme is replaced with a phenylalanine. Thus, without the *cbb*_*3*_-type oxidase found only in *N. gracilis*, it remains unclear how the final electron transport step from cyt. *c* to O_2_ occurs in “*Ca*. N. vancouverensis” and “*Ca*. N. litoralis”. Future biochemical and protein structural research may reveal whether the cyt. *bd*-like oxidases can catalyze this reaction despite their divergence from *bona fide* HCOs at two of the predicted cofactor-binding residues, and whether these proteins are capable of proton translocation for proton motive force generation. Corroborating evidence for a function of this enzyme in the context of electron transport stems from its highly conserved genetic synteny within Nitrospinae, Nitrospirae and anammox bacterial genomes. A conserved cluster within these organisms contains a cyt. *bd*-like oxidase with the aforementioned glutamine and phenylalanine residues, a diheme- and a triheme- cyt. *c*, and a membrane integral, alternative NxrC subunit (Tab. S4). This putative NxrC might be involved in the electron transfer from NO_2_^-^ to the terminal oxidase [25, 63, 66].

In addition to the putative cyt. *bd*-like oxidase discussed above, “*Ca*. N. litoralis” possesses a canonical cyt. *bd*-type (quinol) oxidase that is lacking in “*Ca*. N. vancouverensis” and *N. gracilis*. Since quinol oxidases cannot accept electrons from the high-potential donor nitrite, we assume that this oxidase receives electrons from quinol during the degradation of intracellular glycogen or during hydrogen oxidation (see below). The cyt. *bd*-type oxidase may also be involved in oxidative stress defense, as homologous oxidases in other organisms can degrade H_2_O_2_ [67] and protect from dioxygen [68]. Taken together, the diverse repertoire of terminal oxidases may be a key feature of Nitrospinae that contributes to the ecological success of this lineage over a broad range of redox conditions in marine ecosystems.

### Carbon metabolism and alternative energy metabolisms

Like *N. gracilis* and other Nitrospinae [6, 25], the novel strains encode all key genes of the reductive tricarboxylic acid (rTCA) cycle for CO_2_ fixation, including the hallmark enzymes ATP-citrate lyase (ACL), pyruvate:ferredoxin oxidoreductase (POR), and 2- oxogluterate:ferrodoxin oxidoreductase (OGOR). As in *N. gracilis*, all genes required for the oxidative (oTCA) cycle are also present. All three strains can form glycogen as storage compound, which is degraded via glycolysis and the oTCA cycle. Since *N. gracilis* lacks pyruvate kinase, the final step of glycolysis may be catalyzed by pyruvate phosphate dikinase (PPDK) in this organism. In contrast, strains VA and EB possess both pyruvate kinase and PPDK, indicating a strict regulatory separation between glycolysis and gluconeogenesis.

Alternative energy metabolisms such as the oxidation of hydrogen, sulfide or organic carbon compounds have been demonstrated in NOB with representatives from the genera *Nitrospira, Nitrococcus, Nitrolancea*, and *Nitrobacter* [3, 26–28, 69]. Among the three Nitrospinae strains analyzed here, *N. gracilis* has the largest potential to exploit energy sources other than nitrite: its genome harbors a bidirectional type 3b [NiFe] hydrogenase, which could enable aerobic hydrogen utilization, and a sulfite:cyt. *c* oxidoreductase [25]. In addition, *N. gracilis* contains the genes *prpBCD* for 2-methylisocitrate lyase, 2-methylcitrate synthase, and 2-methylcitrate dehydratase, and might thus be able to catabolically degrade propionate via the 2-methylcitrate pathway (Fig. 5). Of these potential alternative energy metabolisms, “*Ca*. N. litoralis” shares only the type 3b hydrogenase, whereas “*Ca*. N. vancouverensis” seems to be an obligate nitrite oxidizer. No genes for the uptake and utilization of urea and cyanate as organic N sources were found in the genomes of the new strains. These genes show a patchy distribution among Nitrospinae [6, 7, 20, 21, 25], suggesting further niche differentiation of these organisms based on the capacity to use organic compounds as sources of reduced N for assimilation.

**Figure 5.**
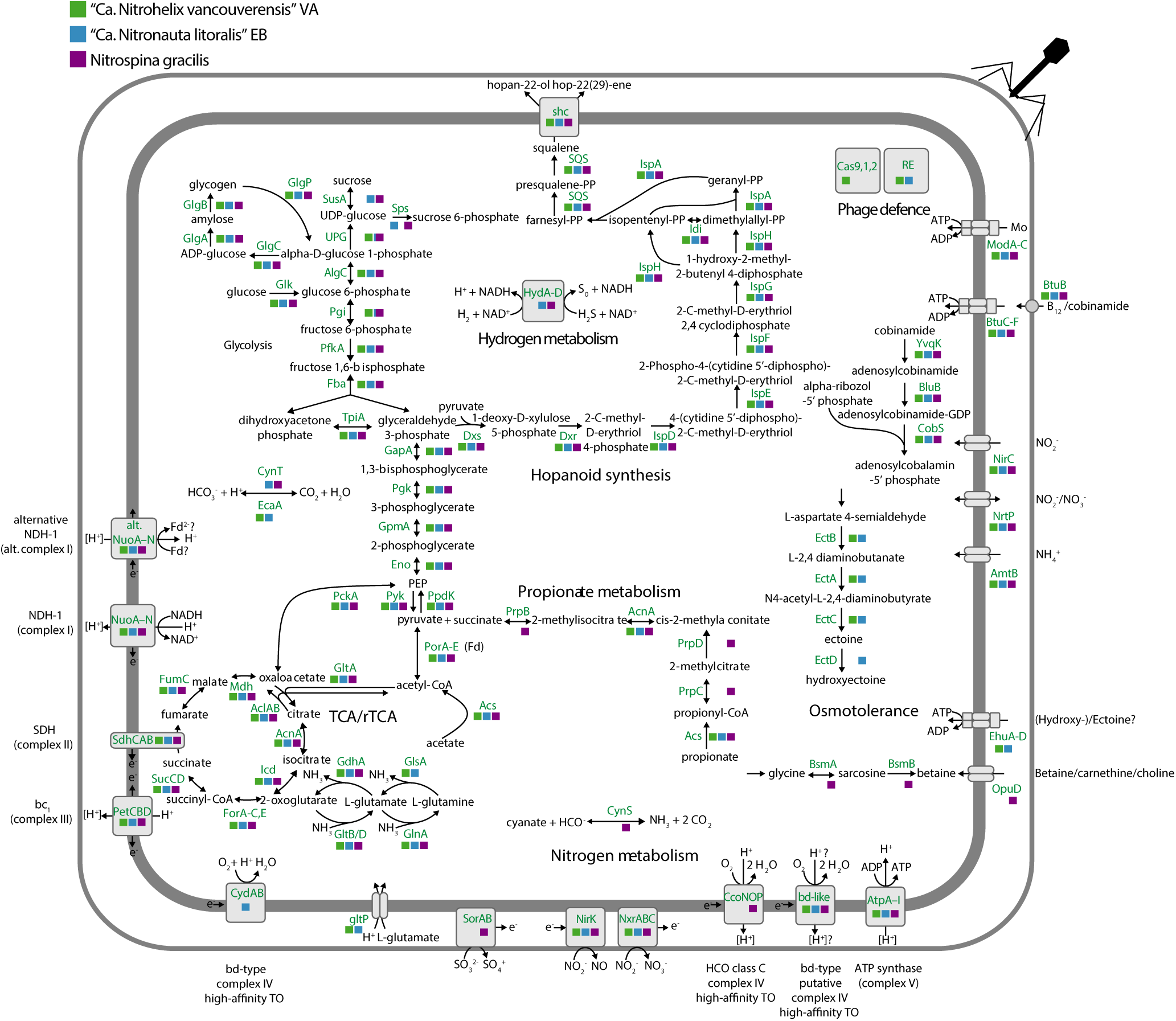
Cell cartoon based on the annotation of “*Ca*. Nitrohelix vancouverensis” VA, “*Ca*. Nitronauta litoralis” EB and *N. gracilis 3/211*. The colored squares indicate the presence or absence of the respective genes, “*Ca*. Nitrohelix vancouverensis” VA is shown in green, “*Ca*. Nitronauta litoralis” EB in blue, and *N. gracilis 3/211* in purple. The gene annotations are detailed in Table S3. Abbreviations: Fd, ferredoxin; RE, restriction enzymes; TO, terminal oxidase.

### Adaptations to saline environments

The intracellular accumulation of ions or the production of organic osmolytes are two main strategies of microorganisms to cope with the osmotic stress in highly saline, marine environments. Interestingly, the three Nitrospinae strains seem to utilize different osmotic stress defense mechanisms. *N. gracilis* has the genetic potential to produce glycine betaine, an organic osmolyte that is ubiquitously found in bacteria (Fig. 5) [70]. It also encodes for OpuD, a betaine/carnitine/choline transporter, whereas the genes for glycine betaine synthesis and import are missing in “*Ca*. N. vancouverensis” and “*Ca*. N. litoralis”. These two strains harbor the canonical genes *ectABC* for ectoine biosynthesis (Fig. 5), another widely distributed osmolyte in bacteria [70]. “*Ca*. N. litoralis” additionally encodes the ectoine hydroxylase EctD and is thus able to form hydroxyectoine. Directly downstream of the (hydroxy-)ectoine synthesis cassette, the two strains further encode an ABC transporter that has similarity to ectoine or amino acid transporters from other organisms and may be utilized for (hydroxy-)ectoine import across the cytoplasmic membrane. Since genes for the synthesis and transport of (hydroxy-)ectoine were also found in “*Ca*. Nitromaritima” (Nitrospinae clade 1) [6], we assume that usage of (hydroxy-)ectoine is wide-spread among the Nitrospinae. *N. gracilis* and “*Ca*. N. litoralis” may also be able to synthesize sucrose as an additional compatible solute (Fig. 5).

Moreover, “*Ca*. N. vancouverensis” and “*Ca*. N. litoralis” genomes harbor the gene *glsA* coding for a glutaminase (Fig. 5), which allows them to deaminate glutamine to glutamate while releasing ammonia. The strains further possess the *gltP* gene coding for a glutamate/H^+^ symporter. Both *glsA* and *gltP* seem to be lacking in the *N. gracilis* genome. Since glutamate can play a role in osmoregulation [70], the ability to regulate the intracellular glutamate level via transport or the degradation of glutamine may be one of various adaptations by “*Ca*. N. vancouverensis” and “*Ca*. N. litoralis” to rapidly respond to stress caused by fluctuating salinities.

All three Nitrospinae strains have the genomic capacity to synthesize the hopanoids hopan-(22)- ol and hop-22(29)-ene (Fig. 5). Hopanoids are pentacyclic, bacterial lipids that integrate into bacterial membranes, much like cholesterol in eukaryotes [71]. They help regulate membrane fluidity and may be important in highly saline environments [70, 72]. Knock-out studies have shown that cells lacking the ability to make hopanoids are more sensitive to various stresses, such as temperature, pH and osmolarity [72]. Interestingly, a metagenomic study of hopanoid- producing bacteria in the Red Sea revealed Nitrospinae to be among the main organisms harboring squalene hopene cyclase, the key gene for hopanoid production [73]. Other marine NOB (members of *Nitrospira, Nitrobacter* and *Nitrococcus mobilis*), and even non-marine *Nitrospira* and *Nitrobacter*, also have the squalene hopene cyclase and therefore the genomic potential to produce hopanoids. Insight into the chemical structure of the hopanoids produced by NOB could help to gain insight into early NOB evolution as hopanoids are important lipid biomarkers and commonly used to deduce ancient microbial activity from sediment fossil records [73].

### Vitamin B12 auxotrophy

The three Nitrospinae strains lack multiple genes involved in vitamin B12 (cobalamin) synthesis and seem to be auxotrophic for this vitamin, as previously suggested for *N. gracilis* [74]. To acquire vitamin B12 from the environment, all strains encode the vitamin B12 ABC transporter *btuCDF* and the outer membrane permease *btuB* [75][74]. Alternatively, this transporter may also import cobinamide that may then be converted to vitamin B12 via a salvage pathway that uses the genes *yvqK, bluB, cobS*, each encoded in all three Nitrospinae genomes [76]. Hence, the availability of externally supplied vitamin B12 or cobinamide is likely of crucial importance for Nitrospinae *in situ* and also in lab cultures. The incomplete cobalamin pathway in the already available *N. gracilis* genome led us to amend the cultivation medium with vitamin B12. Indeed, the addition of vitamin B12, either alone or together with other vitamins, has allowed us to cultivate *N. gracilis* in a more defined medium that is based on Red Sea salt. Previously, the standard medium for this organism had to be prepared from natural seawater [25]. Furthermore, the addition of vitamin B12 was likely an essential prerequisite for our successful enrichment of novel Nitrospinae after cell sorting. The co-cultured alphaproteobacteria may also provide additional vitamin B12 as they both have the genomic repertoire for its synthesis. In the environment, vitamin B12 could be supplied by different heterotrophic or autotrophic microorganisms including ammonia-oxidizing thaumarchaeota, which have been shown to produce vitamin B12 [77–79] and often co-occur with Nitrospinae [10, 21, 80].

## Conclusions

Nitrospinae are important players in marine nitrogen and carbon cycling, but difficulties to cultivate these bacteria have been a major obstacle for their characterization. In this study, the usually very time-consuming enrichment and purification procedure of marine NOB was accelerated by combining cell sorting with genome-informed adjustments to the cultivation medium. By employing this method, we were able to obtain two new, highly enriched Nitrospinae strains, which represent two novel genera in the Nitrospinae and double the number of available cultures from this phylum. A comparison of their completely sequenced genomes and that of *N. gracilis* revealed numerous shared metabolic features, as well as several non-shared, putative adaptations in these distantly related Nitrospinae. With the new cultures at hand, it will now be possible to systematically test such genome-based hypotheses and to elucidate the ecological roles played by members of the Nitrospinae within and beyond the nitrogen cycle.

### Taxonomic consideration of “*Candidatus* Nitrohelix vancouverensis” gen. nov. sp. nov

Ni.tro.he’lix L. n. nitrum: nitrate, L. n. helix: a coil, spiral; N.L. fem. n. *Nitrohelix* nitrate-forming spiral. van.cou.ver.en’sis L. fem. adj. *vancouverensis* of Vancouver.

A nitrate-forming helical bacterium obtained from Vancouver, Canada. Phylogenetically affiliated with the family Nitrospinaceae, phylum Nitrospinae. Cells are gram-negative helically shaped rods with 1-3 turns and a length of approximately 3 µm. The genome consists of a single chromosome of 3,309,797 bp. The DNA G+C content is 51 mol%.

Strain “*Candidatus* Nitrohelix vancouverensis VA” was cultivated from coastal surface sediment from Vancouver, Canada. Marine aerobic chemolithoautotroph that oxidizes nitrite to nitrate. *K*_m(app)_ is 8.7±2.5 µM NO_2_^-^. The strain was routinely cultured with 0.5 mM nitrite at 28°C in liquid marine mineral medium. Could not be grown on solid medium. Auxotrophic for vitamin B12 according to genome analysis.

### Taxonomic consideration of “*Candidatus* Nitronauta litoralis” gen. nov. sp. nov

Ni.tro.nau’ta L. n. nitrum: nitrate, L. n. nauta: seaman; N.L. masc. n. *Nitronauta* nitrate-forming seaman. li.to.ra’lis L. masc. adj. *litoralis* coastal.

A nitrate-forming marine bacterium found in a coastal habitat. Phylogenetically affiliated with the family Nitrospinaceae, phylum Nitrospinae. Cells are gram-negative short rods with a length of approximately 1.5 µm. The genome consists of a single chromosome of 3,921,641 bp. The DNA G+C content is 47 mol%.

Strain “*Candidatus* Nitronauta litoralis EB” was cultivated from coastal surface sediment from Elba, Italy. Marine aerobic chemolithoautotroph that oxidizes nitrite to nitrate. *K*_m(app)_ is 16.2±1.6 µM NO_2_^-^. The strain was routinely cultured with 1 to 5 mM nitrite at 28°C in liquid marine mineral medium. Could not be grown on solid medium. Auxotrophic for vitamin B12 according to genome analysis.

## Supporting information

Supplementary Materials and Methods, Supplementary Results and Discussion, Supplementary Figures S1-S8

## Acknowledgements

We would like to thank Julia Polzin and Jillian Petersen for providing sediment samples from Elba. We are also grateful for the help from Daniela Gruber at the Core Facility of Cell Imaging and Ultrastructure Research at the University of Vienna, who supported sample preparation and visualization of electron microscopy samples. We would further like to thank Karin Kohlweiss for performing FACS and the BOKU Core Facility for *Biomolecular and Cellular Analysis* and EQ-BOKU VIBT GmbH for access to equipment. We greatly appreciate Mads Albertsen and Soeren M. Karst for the Nanopore sequencing conducted during the course “Hands-on Metagenomics using Oxford Nanopore DNA Sequencing”. We also thank Marcel Kuypers, Katharina Kitzinger, Hannah Marchant, and Bela Hausmann for valuable discussions and Bernhard Schink for advice on naming the new organisms. This study was supported by the Austrian Science fund (FWF) project Microbial Nitrogen Cycling – From Single Cells to Ecosystems (W1257). AJM was partially supported by a Fellowship from the Natural Science and Engineering Council of Canada Postgraduate Scholarship-Doctoral (NSERC PGS-D). MYJ was supported by an ERC Advanced Grant project to MW (Nitricare; 294343).

