## Supplementary Materials and Methods, Supplementary Results and Discussion, Supplementary Figures S1-S8 for "Genomic and kinetic analysis of novel Nitrospinae enriched by cell sorting"

#### Isolation of accompanying heterotrophs

Alphaproteobacterial heterotrophs were isolated from Nitrospinae containing co-cultures on Difco Marine Broth Agar 2216 (Difco, BD) at 28°C. Colonies were observed after ~3 days. After colony picking, the alphaproteobacterial strains were grown in liquid Difco Marine Broth (DB) media at 28°C for all subsequent experiments. To identify these organisms, their 16S rRNA genes were PCR-amplified, using the universal bacterial 16S rRNA gene-targeted primer pair 616V-1492R [1], and Sanger-sequenced (Microsynth). The obtained 16S rRNA gene sequences were subjected to BLASTN searches against the NCBI bacterial and archaeal 16S rRNA sequence database. The 16S rRNA gene sequences are available under the GenBank accession numbers MT338542 (*Stappia sp.)* and MT338538 (*Maritimibacter sp.*).

#### Growth curves and nitrite oxidation

Cells from 25 ml of nitrite-depleted cultures were harvested by centrifugation (4500×g, 20 min, 28°C using a swing-bucket rotor). The cells were subsequently washed with 2.5 ml and suspended in 500 µl of nitrite free medium. 100 µl of the washed and concentrated cells were inoculated into 25 ml of the strain-specific marine mineral medium (see main text Materials and Methods) with 1.1 mM and 0.55 mM nitrite for the EB and VA strain, respectively. The experiments were carried out with biological quadruplicates. Samples for quantitative PCR (qPCR) and nitrite/nitrate quantification were regularly taken and stored at -20°C until further analysis.

Standards for qPCR were prepared by amplifying 16S rRNA gene fragments (201 to221 bp) and cloning the resulting PCR products into a pCR-TOPO TA vector (Invitrogen) according to the manufacturer's protocol. The standards were then amplified from the TOPO construct using the M13 primer pair and a high fidelity Q5 polymerase (NEB), before the background plasmid was digested with DpnI (NEB). The product was purified with a PCR clean-up kit (Jena Bioscience) and quantified using the Qbit dsDNA BR kit (Invitrogen). Details on primer sequences and annealing temperatures can be found in Table S5. Copy numbers of the purified, linearized standards were calculated online with the DNA Copy Number and Dilution Calculator (ThermoFisherScientific).

The samples from the growth experiments were diluted 1:10, freeze-thawed at 65°C for three times, and analyzed by qPCR, using the above described standards and primers, the iQ SYBR Green Supermix (Biorad), and a C1000 CFX96 Real-Time PCR system (Biorad). For the qPCR reactions, 5 and 2.5 µl of the diluted samples was added for VA and EB, respectively, resulting in a 20 µl total reaction volume. To ensure cell lysis, a 15 min 95°C step was added at the beginning of the qPCR cycling protocol. In order to estimate the cell numbers of the *Stappia* sp., the obtained 16S rRNA gene copy numbers were divided by two, because the *Stappia* sp. possesses two 16S rRNA gene copies per genome. The remaining strains only have one genomic copy of the 16S rRNA gene.Nitrite and nitrate concentrations were measured as described elsewhere [2].

**DNA extraction, genome sequencing, and genome assembly**

For Illumina sequencing, DNA from the enrichment cultures was extracted by following a phenol/chloroform/isoamyl alcohol-based extraction protocol (for the full protocol see reference [3]). In short, biomass was harvested from 50 ml of the Elba and Vancouver enrichment by centrifugation (4500×g, 20 min, 4°C using a swing-bucket rotor) and stored at -20°C until DNA extraction. Cells were lysed in lysis matrix E tubes (MP Biomedical) at 4 m/s for 30 s in a mix of phosphate buffer at pH 8, TNS buffer with 10% (w/v) SDS and saturated phenol. After extraction with phenol/chloroform/isoamyl alcohol and chloroform/isoamyl alcohol, the DNA was precipitated with 20% (w/v) polyethylene glycol, washed in ice-cold 75% (v/v) ethanol, and resuspended in sterile DNA/RNA free water. For Nanopore sequencing, high molecular weight DNA was extracted using the Power Soil kit (Qiagen) according to the manufacturer's protocol with the following modifications: cells from the Elba and Vancouver Nitrospinae enrichments were harvested from 70 and 200 ml of the culture, respectively, by centrifugation as described in the main materials and methods section. The cells were stored at -20°C and thawed at room temperature for DNA extraction. After the addition of buffer C1 and prior to vortexing, an additional freeze-thaw step (30 min at -80°C followed by 10 min at 65°C) was included to improve cell lysis. The optional incubation steps of 5 min at 4°C after the addition of buffers C2 and C3 were performed.

DNA library preparation, quality control, and sequencing were performed at the Vienna Biocenter Sequencing Core Facility NGS Unit (www.viennabiocenter.org/facilities) on an Illumina HiSeq sequencer (HiSeqV4 PE125) to generate 2×125 bp paired-end reads. To obtain long reads for genome scaffolding, Nanopore library preparation was performed with the 1D Genomic DNA by Ligation Sequencing Kit (SQK-LSK108, Oxford Nanopore) following the manufacturer's recommendations. The library was sequenced on a R9.4.1 flow cell with the MinION device (Oxford Nanopore Technologies). Nanopore reads were basecalled using Albacore (v. 2.3.1, Oxford Nanopore Technologies). Hybrid assemblies and quality control of the Illumina reads and the long reads from Nanopore sequencing were conducted using the default settings in Spades (v. 3.12.0) and Unicycler (v. 0.4.6) [4, 5]. Closed genomes were obtained for the Vancouver Nitrospinae strain after assembly with Spades, and for the Elba Nitrospinae strain after assembly with Unicycler. A high-quality draft MAG (99.7% completeness, 0.75% contamination) and a low-quality draft MAG (63.6%.7% completeness, 2.41% contamination) were obtained from the Unicycler assembly by differential coverage binning with mmgenome [6] for the Elba and Vancouver alphaproteobacterial species, respectively. A cursory look into the MAGs confirmed the presence of a catalase gene, sod gene, and vitamin B12 biosynthesis pathway using MAGE and prokka (v. 1.12) [7, 8] for the Elba and Vancouver Alphaprotobacteria, respectively.

The MAGs are accessible on NCBI under the Biosample SAMN14924457 (Elba Alphaproteobacteria) and SAMN14944009 (Vancouver Alphaproteobacteria).

**16S rRNA phylogeny**

For phylogenetic analysis, 16S rRNA gene sequences were downloaded from the SILVA SSU Ref NR database (release 138) [9] with the following criteria: taxonomy, Nitrospinales; sequence length, >1399 nucleotides; sequence quality score, >90; pintail score, >90. The 16S rRNA genes (>1400 bp) from the Nitrospinae genomes from this study were extracted from the genomes with hmmer (v. 3.2.1) [10] using a bacterial 16S rRNA hidden markov model (HMM) and then added to the sequence dataset. In addition, we added two environmental 16S rRNA sequences that were identified by a BLASTN search of the NCBI nr database and shared over 94% percent identity with either EB or VA. All sequences were subsequently aligned by using the web version of the SINA aligner (v. 1.2.11) [11]. From the resulting alignment, a maximum likelihood phylogenetic tree was calculated with IQ-TREE (v. 1.6.2) (model: TIM2e+R4, chosen by automatic model selection, and 1000 ultrafast bootstrap runs) [12–14]. In total 5 16S rRNA gene sequences from Nitrospinae bin 107, RIFCSPLOW02_12_Full_45_22, RIFCSPLOWO2_01_FULL_39_10, *Geobacter metallireducens* and *Desulfococcus multivorans* genomes (Tab. S2) were used as outgroup. FigTree (v. 1.4.4) (http://tree.bio.ed.ac.uk/software/figtree/) was used for phylogenetic tree visualization, and labels were modified with Adobe Illustrator.

**Genome annotation and pan-genome analysis**

The two Nitrospinae genomes were uploaded to the MicroScope platform (MAGE Workflow version: 1.8**)** [7] for coding sequence (CDS) prediction and automatic annotation. The annotations of selected metabolic pathways (Table S3) were manually refined by following the same annotation criteria as outlined in Lücker *et al.* [15]. For those predicted gene products, which had homologs in *N. gracilis* that fulfilled the annotation criteria, we adopted the same annotation as in *N. gracilis* [16]. For CDSs that lacked homologs in *N. gracilis*, the closest hits fulfilling the annotation criteria from the SwissProt or the TrEMBLE databases (as listed in MAGE) were considered (Table S3). To perform pan-genome analyses, the MAGE "Pan/Core-Genome" tool was used with the MICFAM parameters set to 50% amino acid identity and 80% alignment coverage. To examine highly conserved proteins among the Nitrospinae strains, the same tool was used, but with parameters set to 80% amino acid identity and 80% alignment coverage, while considering only the shared core genome.

**Electron and fluorescence microscopy**

Prior to scanning electron microscopy, 10 ml of cells from actively nitrite-oxidizing cultures were fixed in their respective culture medium with glutaraldehyde (final concentration 2.5% v/v) for 1.5 h at room temperature. The cells were then harvested by centrifugation (4500×g, 20 min, 28°C) in Eppendorf swing rotor buckets before 15 µl of the concentrated cells were spotted onto poly-L lysine coated glass slides and dried at 46°C. The slides were washed three times in 0.1 cacodylate and 5 g/l sucrose buffer before being further fixed in a 1% (w/v) osmium tetroxide solution for 40 min. Subsequently, the slides were again washed three times for 10 min in the cacodylate sucrose buffer. These washing steps were followed by an ethanol dehydration series (30%, 50%, 70%, 90%, 96%, 100% v/v), each for 5 min and with two additional washing steps in 100% ethanol. The samples were then critical point dried with 100% (v/v) ethanol using a CPD 300 (Leica) instrument. The dehydrated samples were mounted onto stubs and gold sputter coated with a JFC-2300HR (JOEL). Images were obtained with a JSM-IT300 scanning electron microscope (JOEL). Image brightness and contrast were adjusted using the ImageJ software [17, 18].

Catalyzed reporter deposition fluorescence *in situ* hybridization (CARD-FISH) was conducted as described elsewhere [19] on cells fixed in 1% (v/v) formaldehyde. The Nitrospinae-specific 16S rRNA-targeted oligonucleotide probe Ntspn759 [20], with zero mismatches to the newly enriched strains, was labelled with horseradish peroxidase (HRP) and applied in order to detect the Nitrospinae cells in the enrichments. In short, the cells were immobilized on glass slides in 0.2% (w/v) low-gelling agarose followed by a dehydration in 100% (v/v) ethanol. Endogenous peroxidases were inactivated with 0.01 M HCl for 10 min at room temperature, and the cells were permeabilized with a lysozyme treatment (10 mg/ml, 1 h, 37°C) followed by a short (30 s) incubation in 0.1 M HCl. Hybridization with the HRP-labelled probe was performed with 10%

(v/v) formamide in the hybridization buffer for 3 h at 46°C, followed by a 5 min washing step in washing buffer at 48°C [21]. After HRP equilibration in 1×PBS for 5 min at room temperature, signal amplification was conducted with Cy3-labelled tyramides (New England Nuclear Corporation) for 30 min at 46°C and followed by washing for 5 min in 1×PBS at 48°C. Finally, nucleic acids were stained with SYBR Green for 10 min. The cells were visualized with a Leica TCS SP8X confocal laser scanning microscope. Brightness and contrast of full images were adjusted using ImageJ [17, 18].

**Phylogeny of Nitrospinae NxrB proteins**

NxrB and other genes from the type II DMSO reductase-like family of molybdopterin-binding enzymes were retrieved from all genomes analyzed in this study using hmmer [10] with the Fer4_11 hmm. Additional, relevant amino acid sequences from the type II DMSO reductase-like family were retrieved from NCBI (see Figure S7 for accession numbers). The sequences were aligned with MAFFT [22] linsi, and the alignment was trimmed with trimal [23] with the flag -automated1. The alignments were used to calculate a maximum likelihood tree with the IQ-TREE web server [12–14] (model: LG+I+G4 with automatic model selection and ultrafast bootstraps=1,000). Proteins from the type II DMSO reductase-like family that are neither Nxr nor Nar from Nitrospinae bin 107, Nitrospinae AG-538-L21, Nitrospinae LS NOB, Nitrospinae RIF_22 and *Desulfococcus multivorans* were used as an outgroup. FigTree version 1.4.4 (http://tree.bio.ed.ac.uk/software/figtree/) was used for the visualization of the tree.

### Tetranucleotide sequence analysis and genome comparisons

Whole genome tetranucleotide signatures were calculated for both strands for each genome with the biostring package in R, as described elsewhere [24]. For the BLAST genome comparisons (Figure S8), the CDS for each genome were predicted using prodigal (v 2.6.3) and all the CDS from each genome were separately blasted against each other using BLASTP (v2.9.0+). Maximum target sequences and an e-value cut-off were set to 1 and $10^{-3}$, respectively. Hits were then removed that had a percent alignment of less than 40% and a percent identity of less than 30%. Lastly, the average percent identities of reciprocal BLAST hits were plotted by the centroid position of each gene in the genome.

### Supplementary Results and Discussion

#### Mobile genetic elements and phage defense

A genome-wide reciprocal BLASTP analysis revealed that parts of the flexible genome of each Nitrospinae strain are clustered into genomic islands, which mostly consist of hypothetical proteins (Fig. S8). One of the genomic islands in "*Ca.* N. litoralis" also encodes a putative iron permease, which may be beneficial since iron is an essential cofactor of NXR and other key enzymes of Nitrospinae. The large genome of "*Ca.* N. litoralis" also contains numerous transposases (Table S2), indicating that genomic rearrangements and lateral gene transfer might have played an important role in the evolution of this organism. Furthermore, the variable genomic regions of "*Ca.* N. litoralis" contain numerous putatively phage-related genes and one type II restriction modification system. In "*Ca.* N. vancouverensis", the variable regions encode three CRISPR-Cas proteins (Cas9, Cas1, and Cas2), which are sufficient for CRISPR-based phage defense [25], and two restriction modification systems of type I and II, respectively. The presence of phage remnants and phage defense mechanisms strongly suggest that "*Ca.* N. vancouverensis" and "*Ca.* N. litoralis" interact with bacteriophages in their natural habitats. This would be consistent with the recent hypothesis that Nitrospinae may be targets of extensive predation, which counteracts their rapid *in situ* duplication rates and could explain the lower abundance of Nitrospinae compared to slower growing AOA in the same ecosystems [20]. It is tempting to speculate that the lack of known phage defense mechanisms in the *N. gracilis* genome might be a consequence of gene loss during three decades of cultivation in the lab after this strain was isolated from seawater [26].

16. Lücker S, Nowka B, Rattei T, Spieck E, Daims H. The genome of *Nitrospina gracilis*

illuminates the metabolism and evolution of the major marine nitrite oxidizer. Front Microbiol. 2013;4:1–19.

24. R Core Team. R: A language and environment for statistical computing. 2019. R Foundation for Statistical Computing, Vienna, Austria.

25. Chylinski K, Le Rhun A, Charpentier E. The tracrRNA and Cas9 families of type II

CRISPR-Cas immunity systems. RNA Biol. 2013;10:726–37.

26. Watson SW, Waterbury JB. Characteristics of Two Marine Nitrite Oxidizing Bacteria , *Nitrospina gracilis nov. ten. nov. sp.* and *Nitrococcus mobilis nov. ten. nov. sp.**. Arch Mikrobiol. 1971;77:203–30.

27. Ngugi DK, Blom J, Stepanauskas R, Stingl U. Diversification and niche adaptations of - *Nitrospina* like bacteria in the polyextreme interfaces of Red Sea brines. ISME J. 2015;10:1–17.

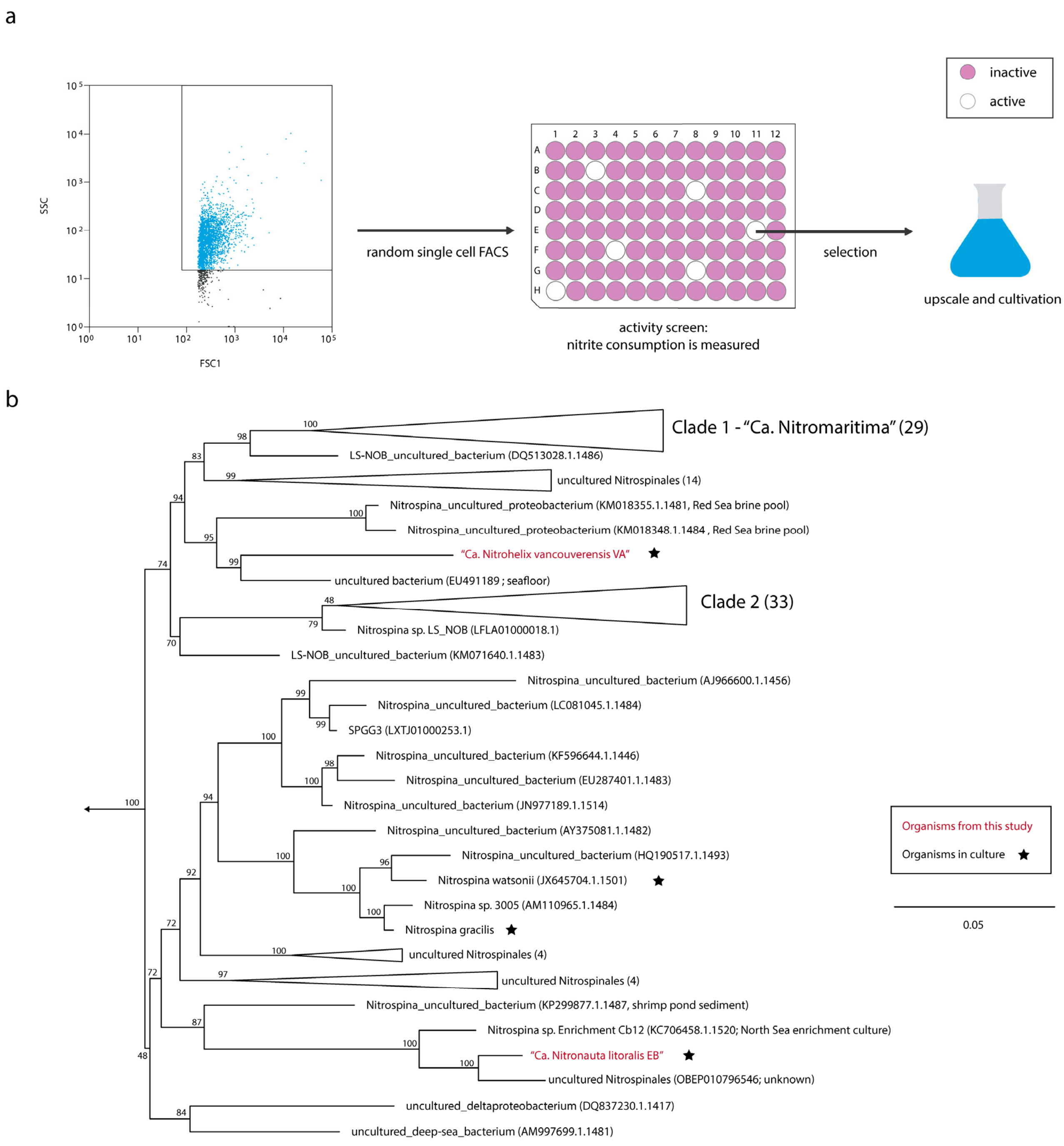


**Figure S1.** Physical enrichment and phylogenetic affiliation of new Nitrospinae. **a** Schematic illustration of the approach to highly enrich novel Nitrospinae strains. Microbial cells were randomly sorted into microtiter plates by fluorescence-activated cell sorting (FACS).

Subsequently, the plates were screened for nitrite-consuming activity by the colorimetric Griess assay. Cells from positive wells were sub-cultured and scaled up. See Materials and Methods in the main text for details. **b** Maximum likelihood tree of selected 16S rRNA gene sequences from the phylum Nitrospinae. Numbers at branches indicate ultrafast bootstrap (n=1,000) support. *D. multivorans* and *G. metallireducens* and the Nitrospinae MAGs bin 107, RIFCSPLOW02_12_Full_45_22, and RIFCSPLOWO2_01_FULL_39_10 were chosen as outgroups. Cultured organisms are marked with asterisk, and the newly cultured strains from this study are highlighted in red. Accession numbers of individual sequences are provided in parentheses and the sample source is indicated for those sequences that group with VA and EB. For collapsed branches (wedges), the numbers of sequences in the respective lineages are indicated in parentheses. The scale bar indicates 0.05 estimated substitutions per nucleotide.

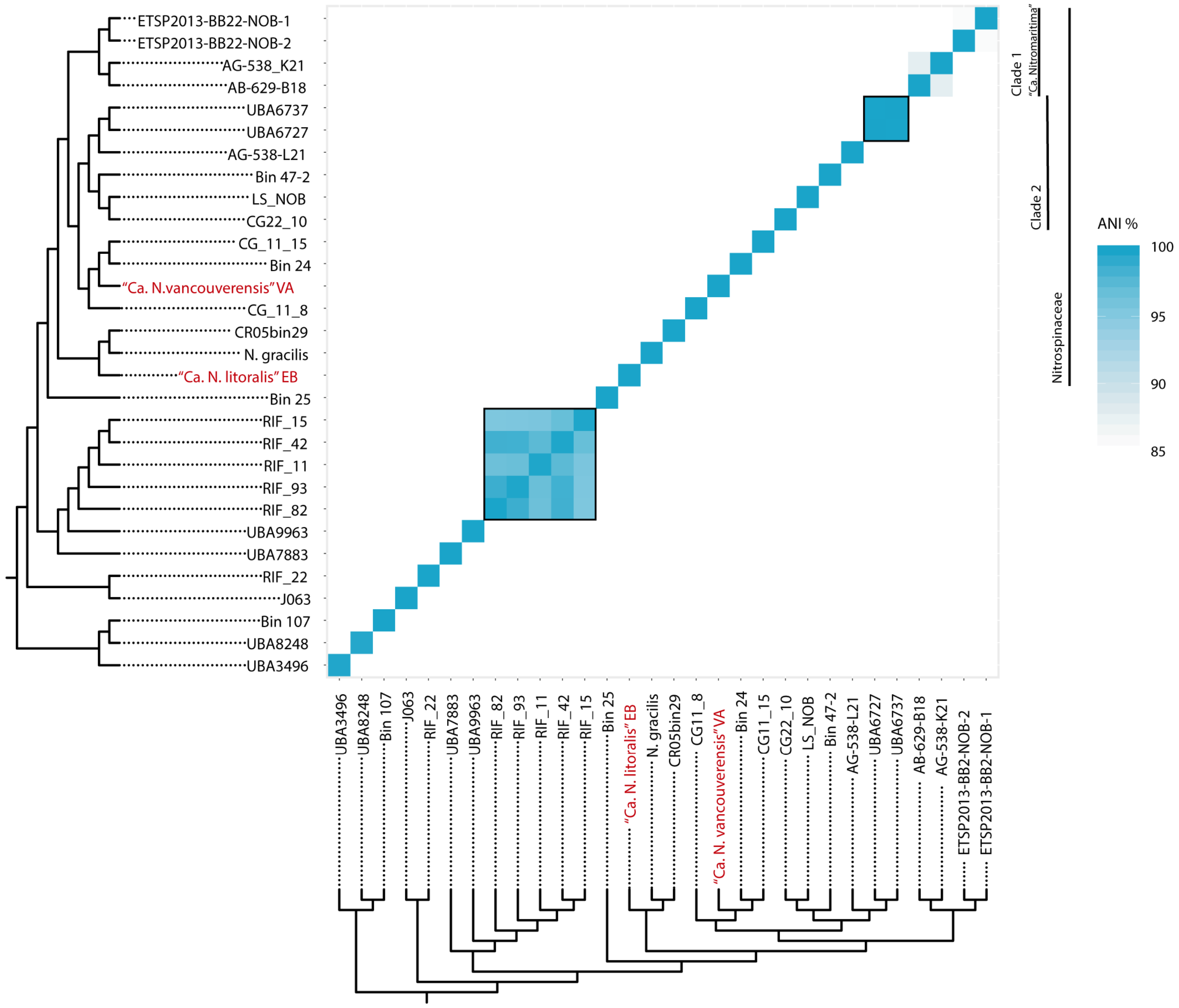


**Figure S2.** Whole-genome average nucleotide identity (gANI) analysis of the Nitrospinae. Pairwise gANI values were calculated for the same set of Nitrospinae genomes that was used to reconstruct the phylogenetic tree in Fig 1 in the main text. The same tree is used here to annotate the heatmap. The two newly cultured strains are highlighted in red. Clades of the Nitrospinae are indicated as proposed elsewhere [27]. Boxes indicate species boundaries with a threshold of 96.5% identity.

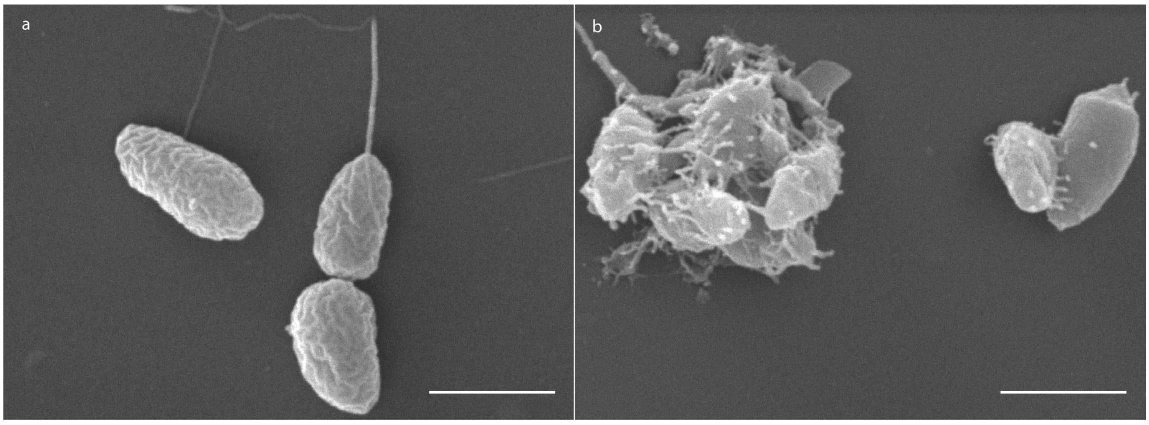


**Figure S3.** SEM images of heterotrophic Alphaproteobacteria co-cultured with the Nitrospinae strains. **a** *Stappia* sp. isolated from the co-culture with "*Ca*. Nitrohelix vancouverensis" strain VA. **b** *Maritimibacter alkaliphilus* isolated from the co-culture with "*Ca*. Nitronauta litoralis" strain EB. The scale bars depict 1 µm.

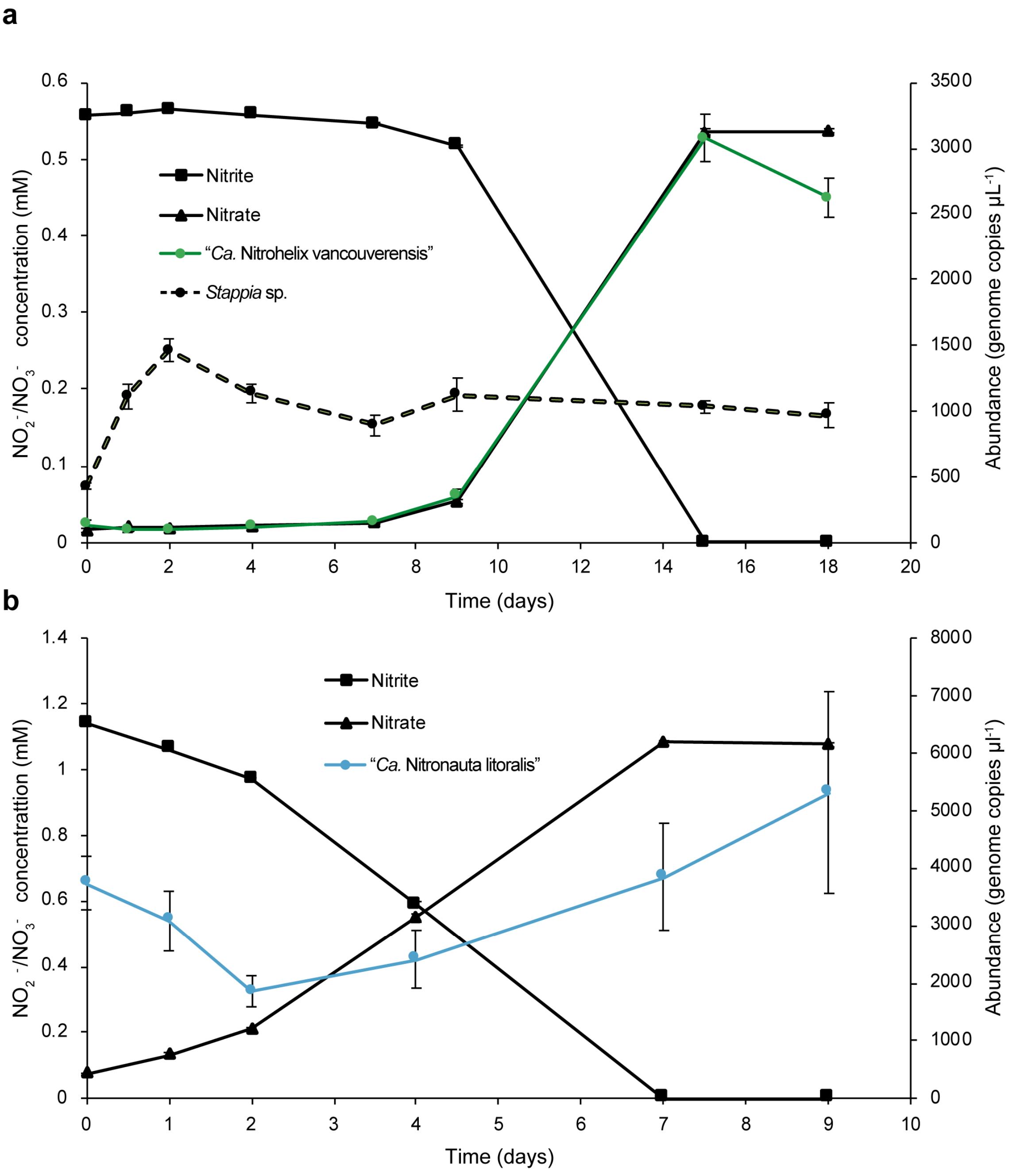


**Figure S4.** Growth and nitrite-oxidizing activity of the new Nitrospinae strains. **a** Growth of "*Ca.* N. vancouverensis" and the co-cultured *Stappia* sp. during consumption of 0.55 mM nitrite. **b** Growth of "*Ca.* N. litoralis" during consumption of 1.1 mM nitrite (the accompanying heterotroph could not be quantified in this experiment). **a, b** Nitrite/Nitrate concentrations are depicted on the left y-axis, and cell abundances measured as genome copies µl$^{-1}$ are shown on the right y-axis. Error bars show the standard error of the mean, based on biological quadruplicates and technical duplicates.

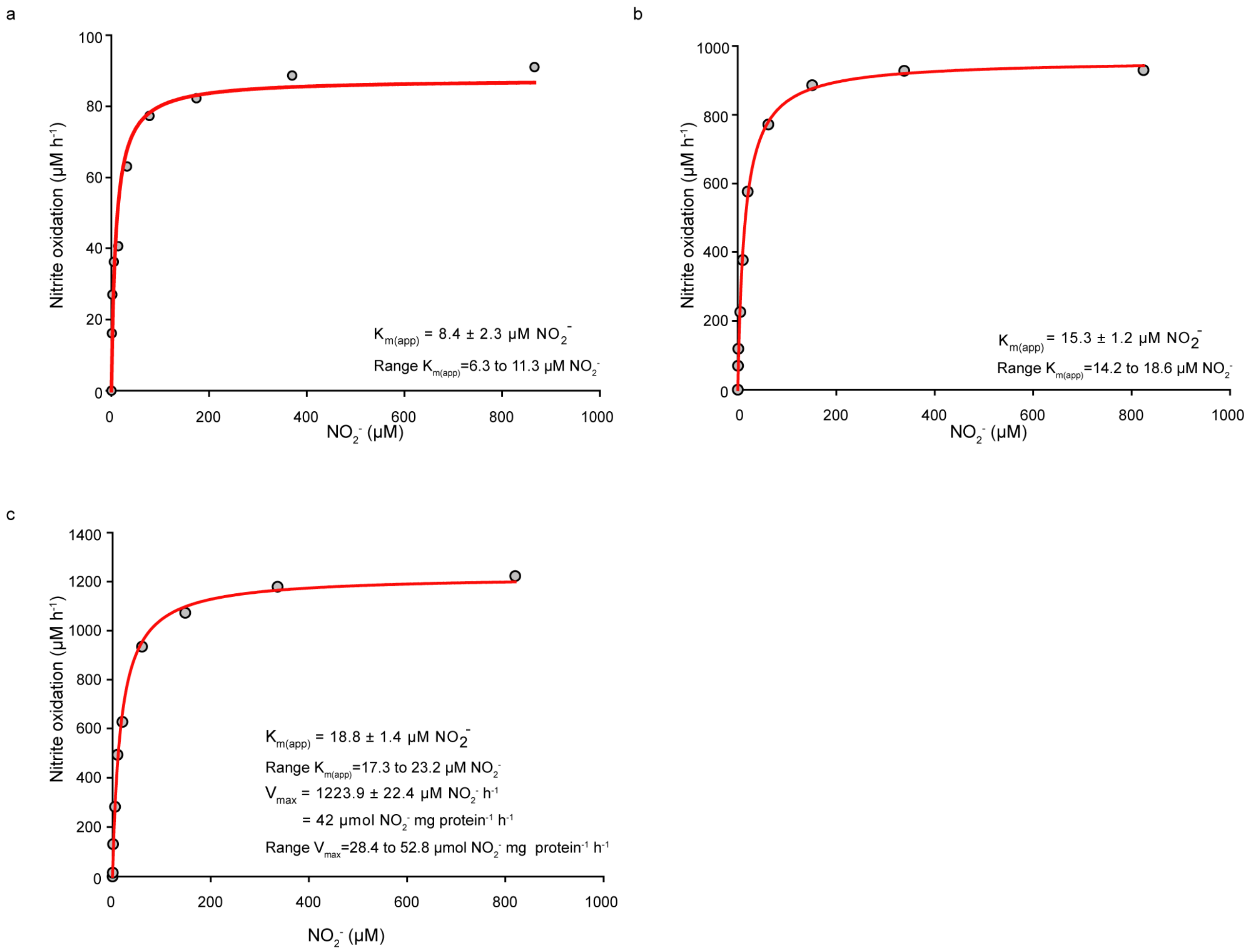


**Figure S5.** Nitrite oxidation kinetics of three Nitrospinae strains. **a-c** Representative Michaelis-Menten plots for "*Ca*. Nitrohelix vancouverensis" strain VA (**a**), "*Ca*. Nitronauta litoralis" strain EB (**b**), and *N. gracilis* (**c**). Nitrite oxidation rates were inferred from nitrite-dependent oxygen consumption. Apparent half-saturation constants, $K_{m(app)}$, and maximum oxidation rates, $V_{max}$, were determined by fitting the data to the Michaelis–Menten kinetic equation. The red curve indicates the best fit of the data. Standard errors of the estimates based on nonlinear regression are reported. Results for one biological replicate are shown; the $K_{m(app)}$ values inferred from all replicates (n=3 for strain VA; n=5 for strain EB and n=8 for *N. gracilis*) are summarized in Fig. 4 of the main text. It should be noted that $V_{max}$ cannot be compared between the three NOB. This would require normalization to biomass (e.g., total protein), which was not possible due to the presence of the co-cultured heterotrophs in the cultures of strains VA and EB.

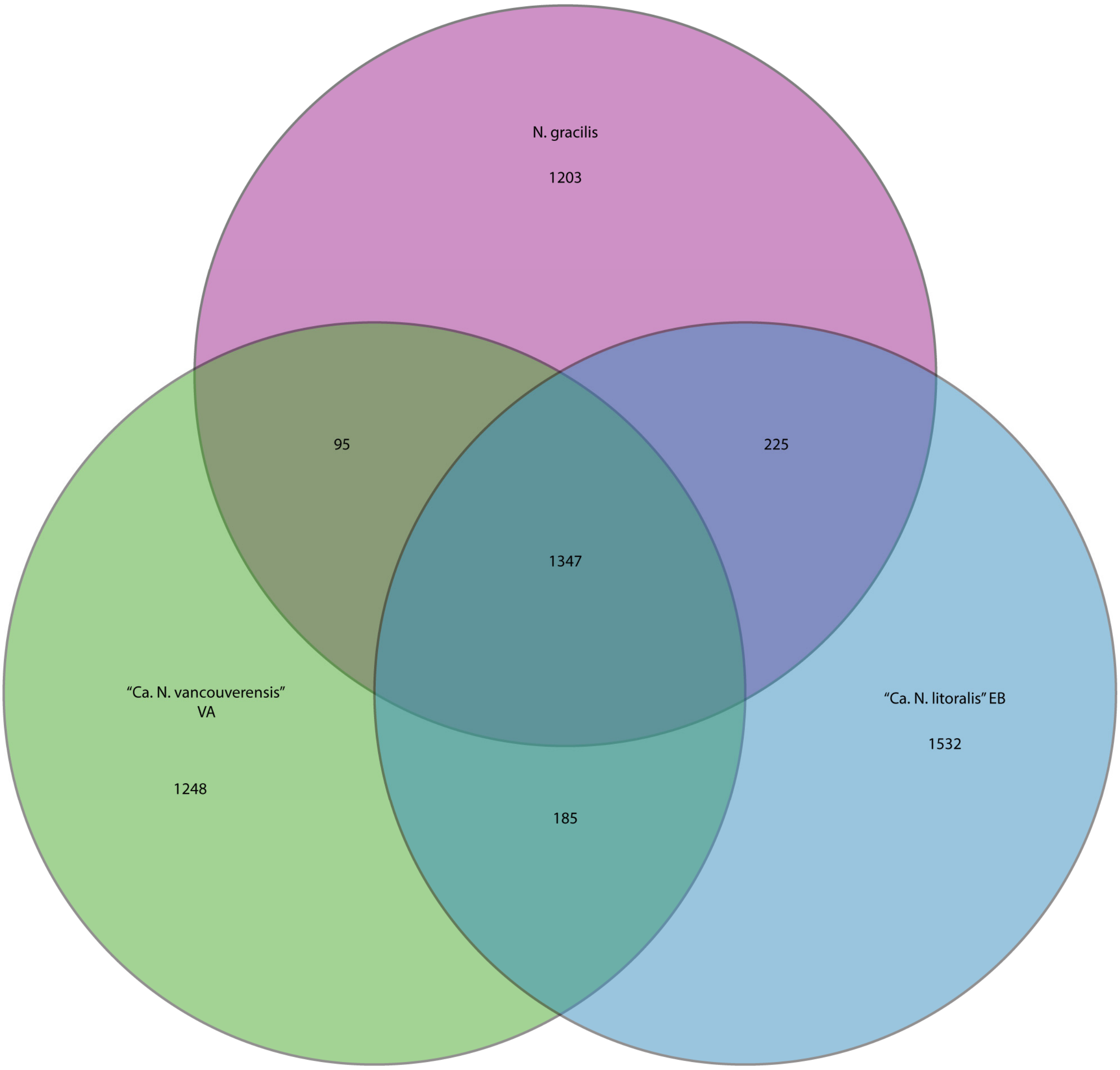


**Figure S6.** Pan-genome analysis of the cultured and genome-sequenced Nitrospinae strains. The Venn diagram shows the numbers of homologous CDS in the shared (core) and variable genomes for *N. gracilis* and the Nitrospinae strains VA and EB. The comparisons were performed by using the MicroScope “Pan/Core-Genome” tool (see Supplementary Materials and Methods).

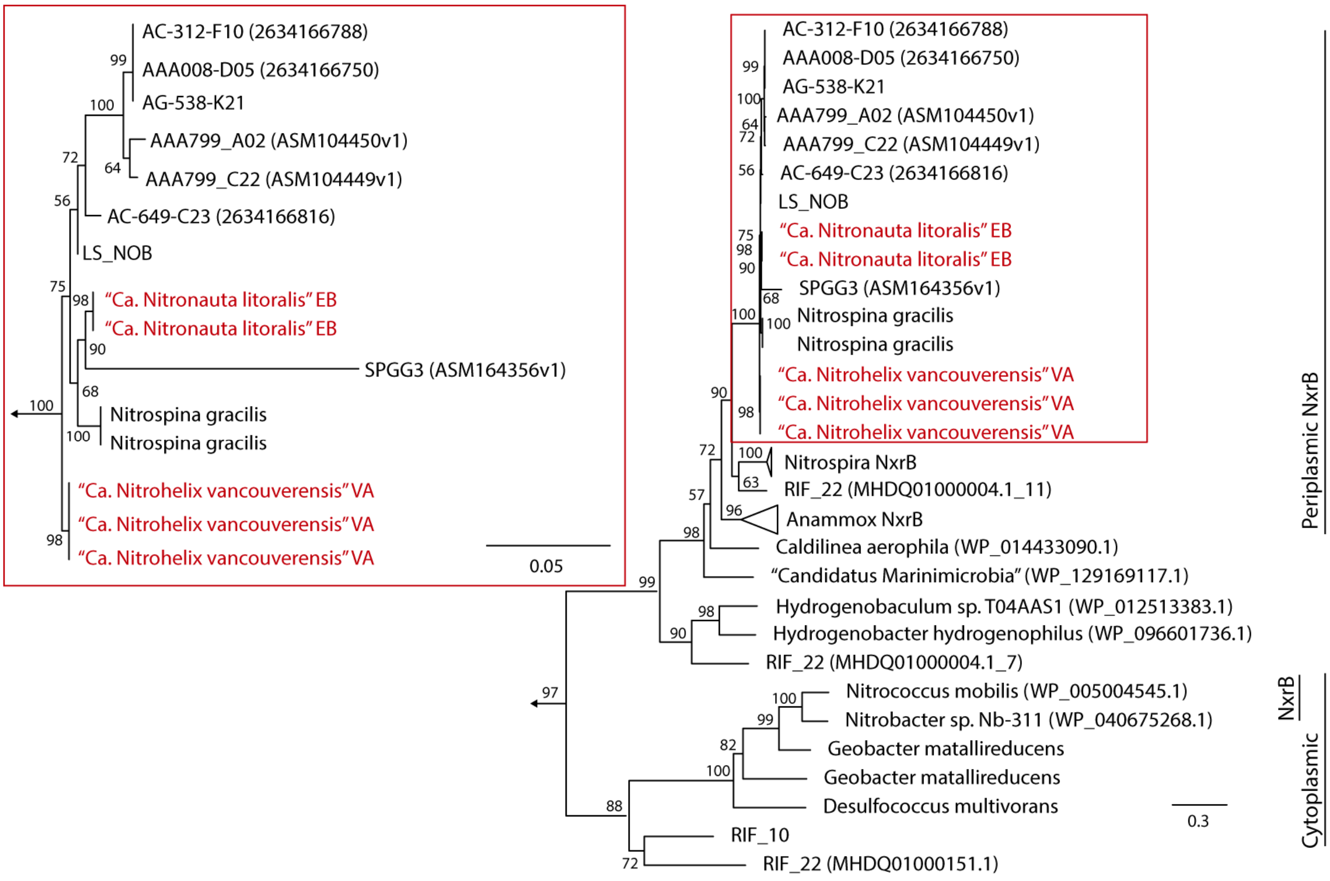


**Figure S7**. NxrB phylogenetic tree. Maximum likelihood tree of NxrB amino acid sequences and other proteins from the type II DMSO reductase-like family of molybdopterin-binding enzymes. Type II DMSO reductase-like family proteins that are neither NxrB nor Nar from Nitrospinae bin 107, Nitrospinae AG-538-L21, Nitrospinae LS NOB, Nitrospinae RIF_22 and *Desulfococcus multivorans* were used as outgroup. The numbers in brackets are the NCBI accession numbers. Organism names in red are the strains obtained in this study. Numbers at branches indicate ultrafast bootstrap (n=1,000) support. The Nitrospinae lineage (red frame) has been enlarged for better visualization of the branching patterns in this group. The scale bars indicate 0.3 and 0.05 estimated substitutions per residue.

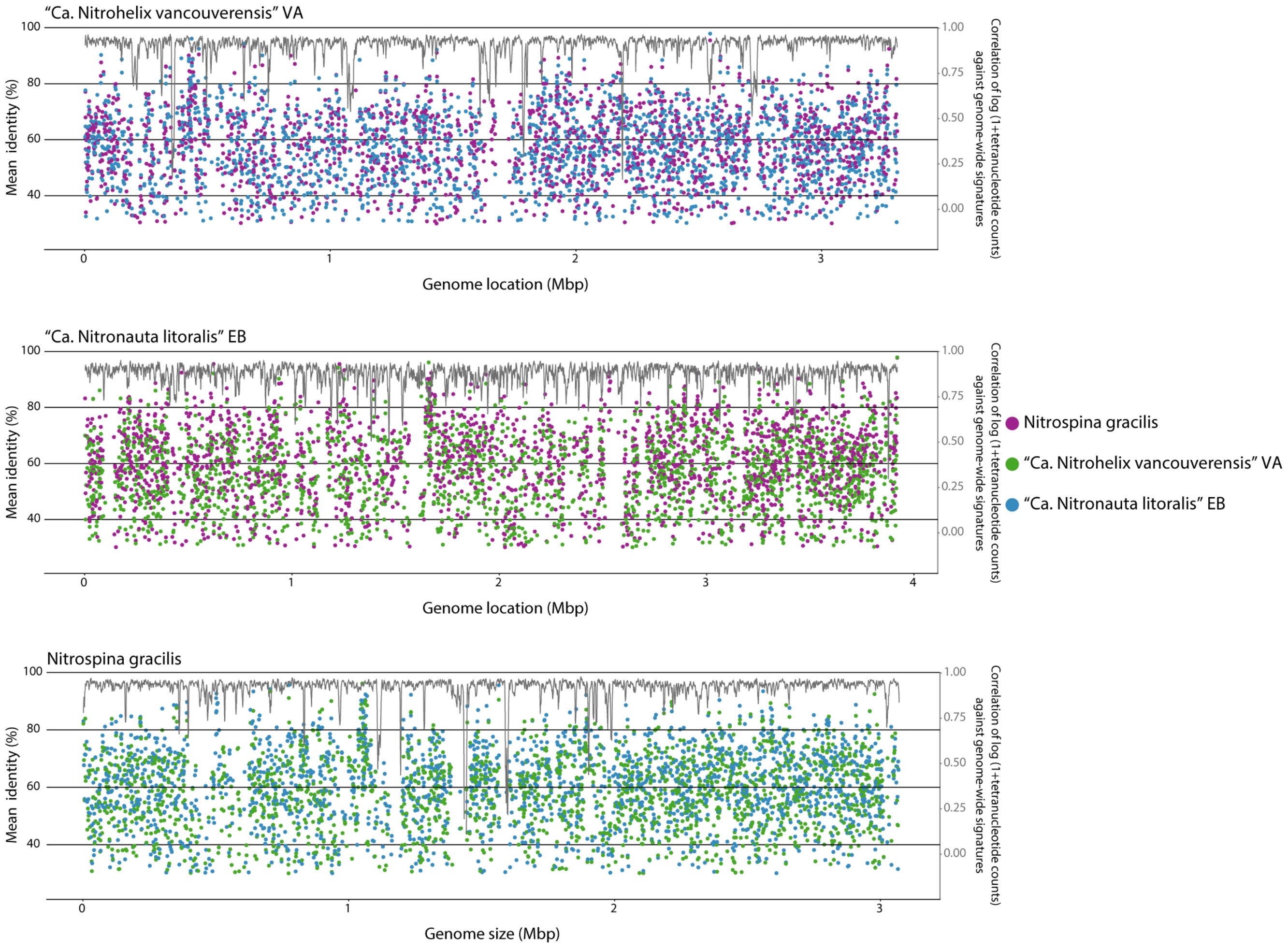


**Figure S8.** Comparative genomics of "*Ca*. Nitrohelix vancouverensis", "*Ca.* Nitronauta litoralis", and *N. gracilis*. Each dot represents a reciprocal BLASTP hit between two of the Nitrospinae strains, the location of dots reflecting the location of the respective CDS in the reference strain (x-axis) and the mean sequence identity (left y-axis). Purple dots refer to *Nitrospina gracilis*, green dots to "*Ca.* Nitrohelix vancouverensis" VA and blue dots to "*Ca.* Nitronauta litoralis" EB. The right y-axis and the grey line show the correlation of the tetranucleotide patterns across a genome, calculated with a 5 kb sliding window and step size of 1 kb, against the genome-wide tetranucleotide signatures.
